# Quantifying Electrostatic Control of Docking and Binding Energetics in Functional Cx36 Gap Junctions

**DOI:** 10.1101/2025.10.25.684567

**Authors:** Robert S. Wong, Zhiyuan Song, Yu T. Zheng, Honghong Chen, Haiqing Zhao, Donglin Bai

## Abstract

Connexin36 (Cx36) is broadly expressed in neurons and serves as the principal protein that forms interneuronal gap junctions (GJs), also known as electrical synapses. Recent high-resolution structures of human Cx36 GJ have revealed crucial electrostatic interactions (ESIs) of charged residues between two docked Cx36 hemichannels at the second extracellular (E2) loops. Despite their structural importance, the mechanistic roles of these ESIs remain poorly understood. To investigate their significance, we systematically designed and tested a series of missense variants targeting key E2 interface residues, aiming to disrupt or modulate the electrostatic landscape at the docking interface. Based on the ESI pairs defined from the crystal structure, our combined computational calculations and dual patch-clamp experiments in engineered HEK293 cell pairs suggest that at least three ESI residual pairs per E2-E2 interface are required to support functional GJ formation. Furthermore, we found that these unique ESIs of Cx36 could play a role in its docking specificity to itself, as they rarely form heterotypic GJs with other brain connexins. Overall, these findings provide essential molecular and functional insights into the mechanisms governing Cx36 GJ formation and partner specificity, paving the way for future therapeutic approaches targeting connexin dysfunction in human diseases.

## Introduction

Direct intercellular communication is essential for coordinating physiological processes within tissues/organs of the body. In the central nervous system, gap junctions (GJs, also known as electrical synapses) serve as specialized structures that enable electrical and metabolic coupling between adjacent neurons in many brain regions^1, 2, 3^. GJs are composed of connexin proteins, which are encoded by 21 distinct genes in the human genome, with each gene encoding one different connexin. Dysfunction or dysregulation of connexin-mediated GJ communication in the brain has been implicated in numerous human diseases, including epilepsy^4^, neurodegenerative diseases^5^ (e.g., Alzheimer’s and Parkinson’s diseases), and autism spectrum disorders^6^, highlighting their critical physiological importance.

From a structural perspective, all connexins share the same topological structure with four transmembrane domains (M1-4), two extracellular loops (E1 and E2), one cytoplasmic loop (CL), and both the amino terminus (NT) and carboxyl terminus (CT) on the cytosolic side^2, 7^. Six connexins oligomerize to form a hemichannel, and two opposing hemichannels dock head-to-head at their extracellular domains to form a GJ channel. When both hemichannels are composed of the same connexin, the resulting GJ is termed homomeric homotypic. If the hemichannels differ, the GJ is either homomeric heterotypic (identical connexins within each hemichannel but different across the junction) or heteromeric heterotypic (different connexins within and across hemichannels). Multiple connexins are often co-expressed within a single cell type or tissue, allowing for the formation of heterotypic GJs not only between cells of the same type, but also at the cell-cell junctions between different cell types^8, 9^. The docking compatibility of different connexins has been found to be selective, and only docking compatible connexins can form functional heterotypic GJs^9, 10^.

Cx36 is the main neuronal connexin, forming GJs or electrical synapses between neurons to mediate direct electrical communications^11, 12, 13^. Genetic ablation of the Cx36 gene in mice has led to impaired electrical coupling and loss of gamma oscillations^14, 15^. Cx36 is also expressed in pancreatic β cells, where it acts as an important regulator for insulin secretion^16, 17^. Following the initial cloning of a fish ortholog of Cx36^18^, mouse, rat, and human Cx36 genes have also been cloned^3, 19, 20^. Functional studies on recombinantly expressed mouse Cx36 (mCx36) revealed that mCx36 can form functional homomeric homotypic GJs but is unable to form homomeric heterotypic GJs with several other connexins, including Cx26, Cx30, Cx31, Cx32, Cx37, Cx40, Cx43, and Cx50^21, 22^. Additionally, a morphological study with immunolabeled rodent Cx45 and Cx36 indicated these two connexins were unable to form morphological homomeric heterotypic GJs^23^. The structural basis for this unique mCx36 docking selectivity remains unclear. Several patch clamp studies have revealed some unique channel properties of the mCx36 GJ^24, 25, 26^, while the docking compatibility of human Cx36 (hCx36) GJs remains unknown. Given the high sequence identity of hCx36 with mCx36 (98%), we hypothesized that hCx36 can only dock with hCx36 to form functional homotypic GJs and is unable to form heterotypic GJs with other human brain-expressed connexins.

After the first high-resolution GJ structure of Cx26 GJ^27^, several other GJs (such as Cx32, Cx46, Cx50, and Cx43) were recently resolved with near-atomic resolution^28, 29, 30, 31, 32, 33^. All these GJ structures showed non-covalent interactions at the docking interface between the two hemichannels of a GJ. Specifically, multiple hydrogen bonds (HBs) were identified between the extracellular loops of the docked subunits, where the E1 domain interacts with two other E1 domains, and the E2 domain interacts with one other E2 domain in the opposing hemichannel^10^. Interestingly, at the E1-E1 docking interface, conserved residues are found to form HBs regardless of the connexin type. In contrast, at the E2-E2 docking interface, hydrogen bonding residues only appear conserved for docking compatible connexins (for example, Cx26 and Cx32)^10^. Previous mutagenesis combined with functional studies demonstrated that the HB-forming residues of E2 loop are critical for the successful formation of functional GJ channels and docking specificity^34, 35, 36, 37, 38^. Together, these observations indicate that the E2-E2 docking interaction is a key factor in deciding the connexin docking compatibility or specificity.

In the case of Cx36, the E2-E2 interface of their gap junction channels is predominantly stabilized by the electrostatic interactions (ESIs) between oppositely charged residues (lysine, K, and three glutamate residues, E), different from the HBs observed in most other connexins^39, 40, 41^. It is not clear how these charged residues at the E2 loop determine the docking compatibility and formation of functional hCx36 GJs. Here, we employ a combined computational and experimental approach to design and examine a series of single and multiple missense variants of the ESI-forming residues on hCx36. These variants were designed to eliminate, reduce, or reverse the ESI interactions at the E2-E2 docking interface between cell pairs expressing the same (homotypic) or different (heterotypic) variants. Our findings revealed an important role for ESIs in hCx36 docking and functional GJ formation. Furthermore, the unique docking mechanism of Cx36 is likely to play a role in its docking specificity, as none of the other brain-expressed connexins were able to form functional homomeric heterotypic GJs with Cx36 *in vitro*. Overall, the revealed docking mechanism of Cx36 advances our understanding of a highly specific and finely tuned intercellular communication network that facilitates the precise neuronal communication via Cx36 gap junctions.

## Results

### The essential role of K238 in forming functional GJs

From the Cryo-EM structure of Cx36 GJ, it is clear that each GJ is composed of two head-to-head docked hexameric Cx36 hemichannels, forming six protein–protein interaction interfaces (known as the docking interface), including interactions between the extracellular domains of docked subunits. Previous studies on other GJs demonstrated that E2-E2 docking interactions are the most important for the docking specificity and formation of functional GJs^34, 35^. At each E2-E2 interface, four charged residues (K238/E239/E230/E241) from each Cx36 protein participate in close-range electrostatic interactions (ESIs) with the same residues of the opposing Cx36 (Fig. 1A). To determine whether the electrostatics of these charged residues play the dominant role in mediating binding affinity between docked hemichannels, we computationally introduced mutations to all four charged residues by substituting them with either neutral or oppositely charged amino acids (see Table 1 for mutation details) and calculated the corresponding changes in the hemichannel binding free energy (ΔΔG) in homotypic GJs. Our results demonstrate that changes in electrostatic interaction energy (ΔΔΨ) strongly govern the observed overall ΔΔG values (Fig. 1B), with a Pearson correlation coefficient of 0.90. This correlation is substantially stronger than that of other interaction types, including hydrogen bonds and *van der Waals* forces (Fig. S1), underscoring the central role of electrostatics in stabilizing the docked Cx36 hemichannels. In the following experiments, we altered these charged interface residues and assessed the effects of electrostatic interactions on GJ function.

**Figure 1.**
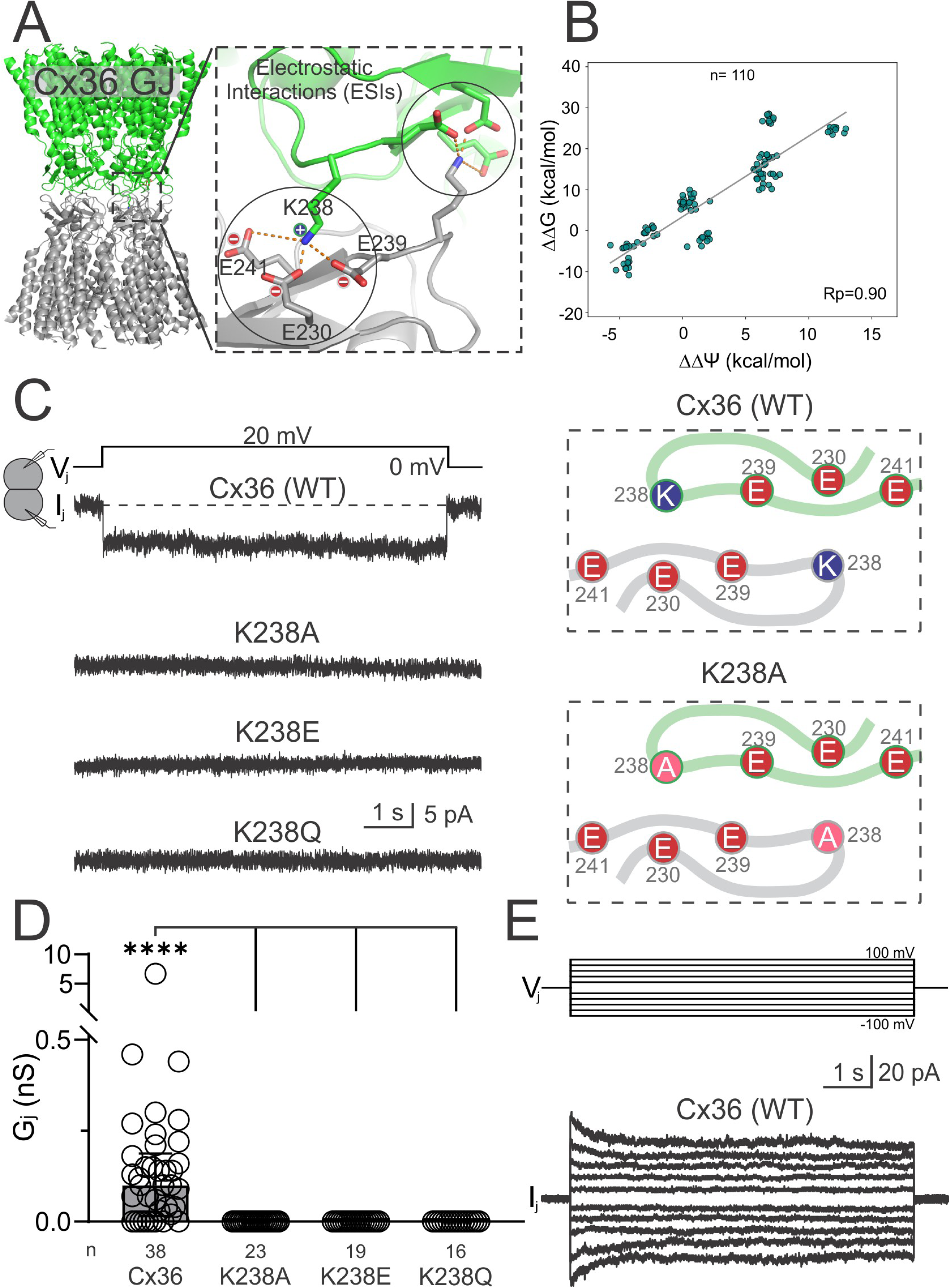
Cx36 K238A, K238E, and K238Q variants cannot form functional GJ channels. (A) Structural model of a Cx36 GJ (PDB: 8XGD) and a zoom-in view of the E2-E2 docking interface. Charged residues and potential electrostatic interactions of the E2-E2 interface are shown on the right. (B) *In silico* calculations on our designed homotypic variant GJs show a strong positive correlation between changes in electrostatics (ΔΔΨ) and the changes in binding free energy (ΔΔG) with a Pearson correlation coefficient of 0.9, exceeding the correlations observed for other energetic components (Fig. S1). Notably, ΔΔΨ accounts for approximately half of the total ΔΔG change. *n* denotes the total number of samples obtained from 10 independent runs of energy calculation across 11 homotypic variants. (C) Representative junctional currents (I_j_s) were recorded in response to a +20 mV transjunctional voltage (V_j_) pulse applied to DKO HEK293 cell pairs expressing the Cx36 variant or wildtype as indicated. A schematic version of Cx36 and a variant (K238A) at the E2-E2 docking interface are shown as an example. (D) Bar graph summarizing the median (error bars represent interquartile range, IQR) junctional conductance (G_j_) for all recorded cell pairs. Statistical significance (\*\*\*\**P* < 0.0001) is obtained from a Kruskal-Wallis test followed by Dunn’s multiple comparisons tests comparing G_j_ of each Cx36 variant to that of wildtype. “n” is the number of biologically independent cell pairs measured for each group. (E) Superimposed I_j_s measured in a Cx36-expressing cell pair, in response to a series of V_j_ pulses ranging from ±20 mV to ±100 mV, at 20 mV increments for 7 s duration each.

**Table 1.**
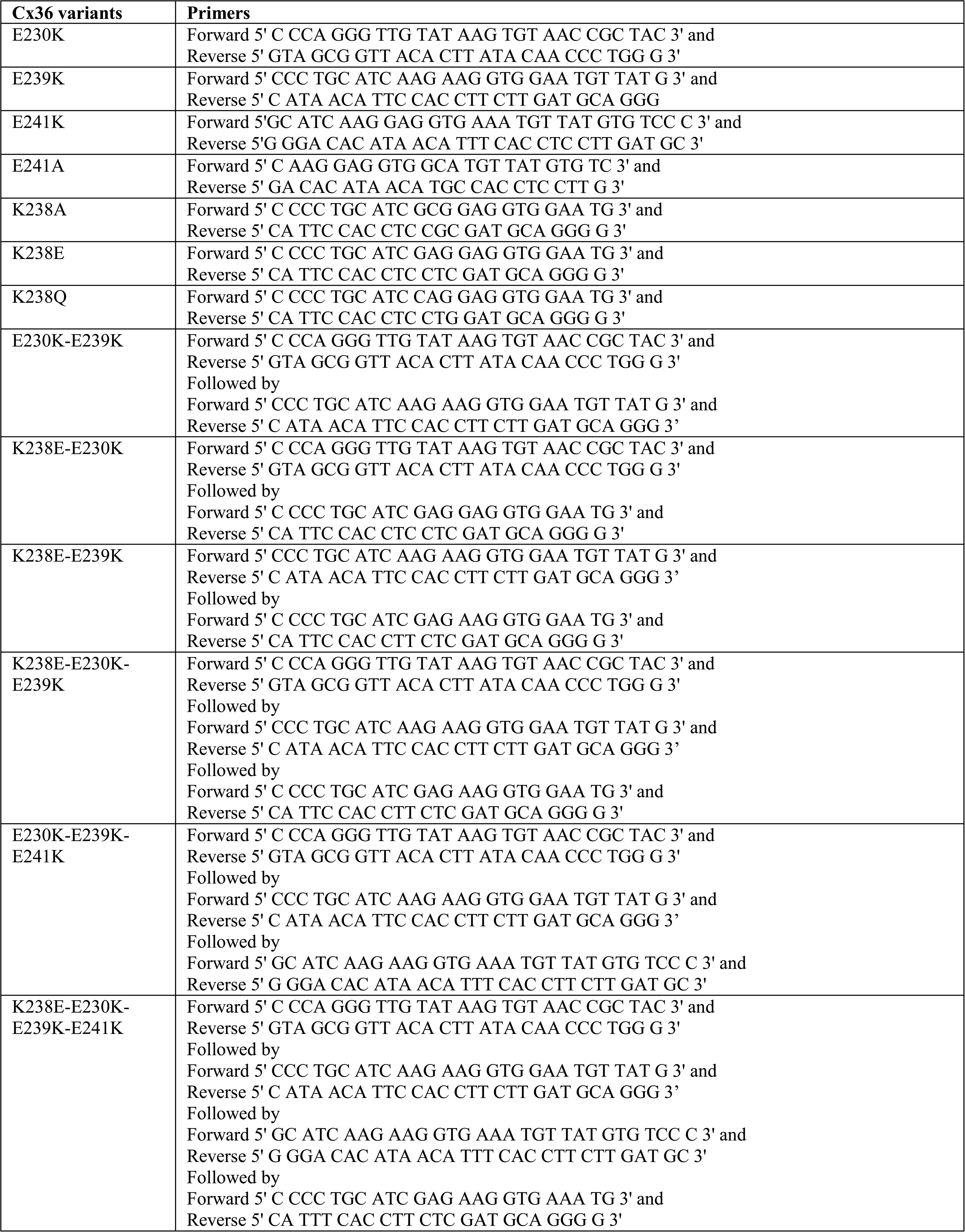
Primers used to generate Cx36 variants.

We performed dual whole cell patch clamp on connexin-deficient HEK293 (Cx43 and Cx45 double knockout HEK293 or DKO HEK293)^42^ cell pairs expressing Cx36 variants in an expression vector with an untagged reporter (i.e. variant-IRES-EGFP or variant-IRES-DsRed). Transjunctional current (I_j_) was measured in response to a +20 mV transjunctional voltage (V_j_) pulse to assess GJ function in the cell pair. Empty expression vector (with IRES-GFP or IRES-DsRed) expressing DKO HEK293 cells were routinely used as a negative control to check for background GJ coupling. To eliminate all E2-E2 ESIs at the hemichannel docking interface, three different Cx36 variants were designed around the key positive interface residue K238 (Fig. 1C): K238A to neutralize the charge of the lysine sidechain, K238E to reverse the positive charge of the lysine sidechain to a negatively charged glutamate, and K238Q, which replaced the positive lysine with a non-charged polar glutamine. All of these variants were predicted to eliminate the ESIs at the E2-E2 docking interface. Although I_j_ is a signature observation for wildtype Cx36-expressing cell pairs, no I_j_ response was observed in any of the designed K238 variants expressing cell pairs (Fig. 1C). The junctional conductance (G_j_) for each tested group was calculated from individual cell pairs and is presented in Fig. 1D, showing the median G_j_ (interquartile range, IQR) for cell pairs recorded (Table 2). K238A, K238E, and K238Q variant-expressing cell pairs showed no GJ coupling, with G_j_s significantly lower than that recorded from Cx36-expressing cells. When Cx36-expressing cell pairs were administered with a series of V_j_ pulses (±20 mV to ±100 mV), the corresponding I_j_s exhibited mirror symmetrical deactivations upon higher V_j_ pulses (±80 to ±100 mV) demonstrating minor V_j_-gating as shown in Fig. 1E.

**Table 2.** Median G_j_s (IQR) for all Cx36 variant combinations.

| **Homo- or hetero-typic $G_j$s†** | **Median $G_j$ (nS)** | **IQR** | **# pairs** | **# trans.** |
| --- | --- | --- | --- | --- |
| Cx36 | 0.1 | 0.02 - 0.19 | 38 | 11 |
| K238A | 0 | 0 - 0 | 23 | 5 |
| K238E | 0 | 0 - 0 | 19 | 5 |
| K238Q | 0 | 0 - 0 | 16 | 4 |
| E230K | 0.01 | 0 - 0.14 | 28 | 7 |
| E239K | 0.12 | 0.04 - 0.31 | 30 | 7 |
| E241K | 0 | 0 - 0 | 25 | 6 |
| E230K-E239K | 0 | 0 - 0 | 20 | 4 |
| K238E-E230K-E239K-E241K | 0 | 0 - 0 | 16 | 4 |
| K238E-E230K-E239K | 0.12 | 0.08 - 0.14 | 16 | 4 |
| K238E-E230K | 0 | 0 - 0 | 15 | 4 |
| K238E-E239K | 0 | 0 - 0 | 15 | 4 |
| K238E / E230K-E239K-E241K | 0 | 0 - 0 | 18 | 4 |
| K238E / E241K | 0 | 0 - 0 | 19 | 5 |
| E230K-E239K-E241K | 0 | 0 - 0 | 12 | 3 |
| E241A | 0.09 | 0.03 - 0.19 | 18 | 5 |
| Cx36 / E230K | 0.14 | 0.04 - 0.22 | 15 | 3 |
| Cx36 / E239K | 0.13 | 0.08 - 0.19 | 18 | 3 |
| Cx36 / E230K-E239K | 0.22 | 0.11 - 0.41 | 19 | 3 |
| E230K-E239K / K238E-E230K | 0.09 | 0 - 0.18 | 15 | 3 |
| E230K / K238E | 0.1 | 0.05 - 0.22 | 20 | 5 |
| E239K / K238E | 0.11 | 0.08 - 0.2 | 23 | 5 |
| Cx36 / K238A | 0.16 | 0.05 - 0.24 | 17 | 3 |
| Cx36 / K238Q | 0.13 | 0.01 - 0.31 | 16 | 3 |
| E239K / E230K-E239K | 0.05 | 0 - 0.12 | 21 | 4 |
| Cx36 / K238E-E239K | 0.06 | 0 - 0.10 | 15 | 3 |
| E241A / E230K-E239K | 0 | 0 - 0 | 16 | 3 |
| E230K / E230K-E239K | 0 | 0 - 0 | 20 | 4 |
| Cx36 / K238E-E230K | 0 | 0 - 0 | 16 | 3 |
| K238E / K238E-E230K | 0 | 0 - 0 | 15 | 3 |
| K238E / K238E-E239K | 0 | 0 - 0 | 15 | 3 |

†Homo-typic $G_j$ s are indicated by variant alone, and heterotypic $G_j$ s are indicated by “variant1 / variant2”; “IQR”, interquartile range; “# pairs” represents the number of cell pairs recorded for each combination; “# trans.” represents the number of transfections.

### Designed Cx36 variants with reduced or reversed E2-E2 ESIs

To test the ability of Cx36 variants to form functional GJs with different levels of reduced E2-E2 ESIs (based on counting the predicted number of possible ESIs, see Fig. S2), the following Cx36 variants were designed and tested for their GJ function using dual patch clamp: E230K, E239K, E241K, and E230K-E239K. I_j_s were observed in response to a V_j_ pulse in E230K and E239K variant-expressing cell pairs, but not in cell pairs expressing E241K or E230K-E239K (Fig. 2A). The median G_j_ (IQR) of all cell pairs recorded (Table 2) was summarized in a bar graph (Fig. 2B). The G_j_s of E230K and E239K were not statistically different from that of Cx36, while E241K and E230K-E239K displayed no coupling (***P < 0.001 comparing to Cx36), indicating that reducing the predicted E2-E2 ESIs still resulted in successful formation of functional GJs except E241K. Further reduction of the predicted E2-E2 ESIs with E230K-E239K was unable to form functional GJs.

**Figure 2.**
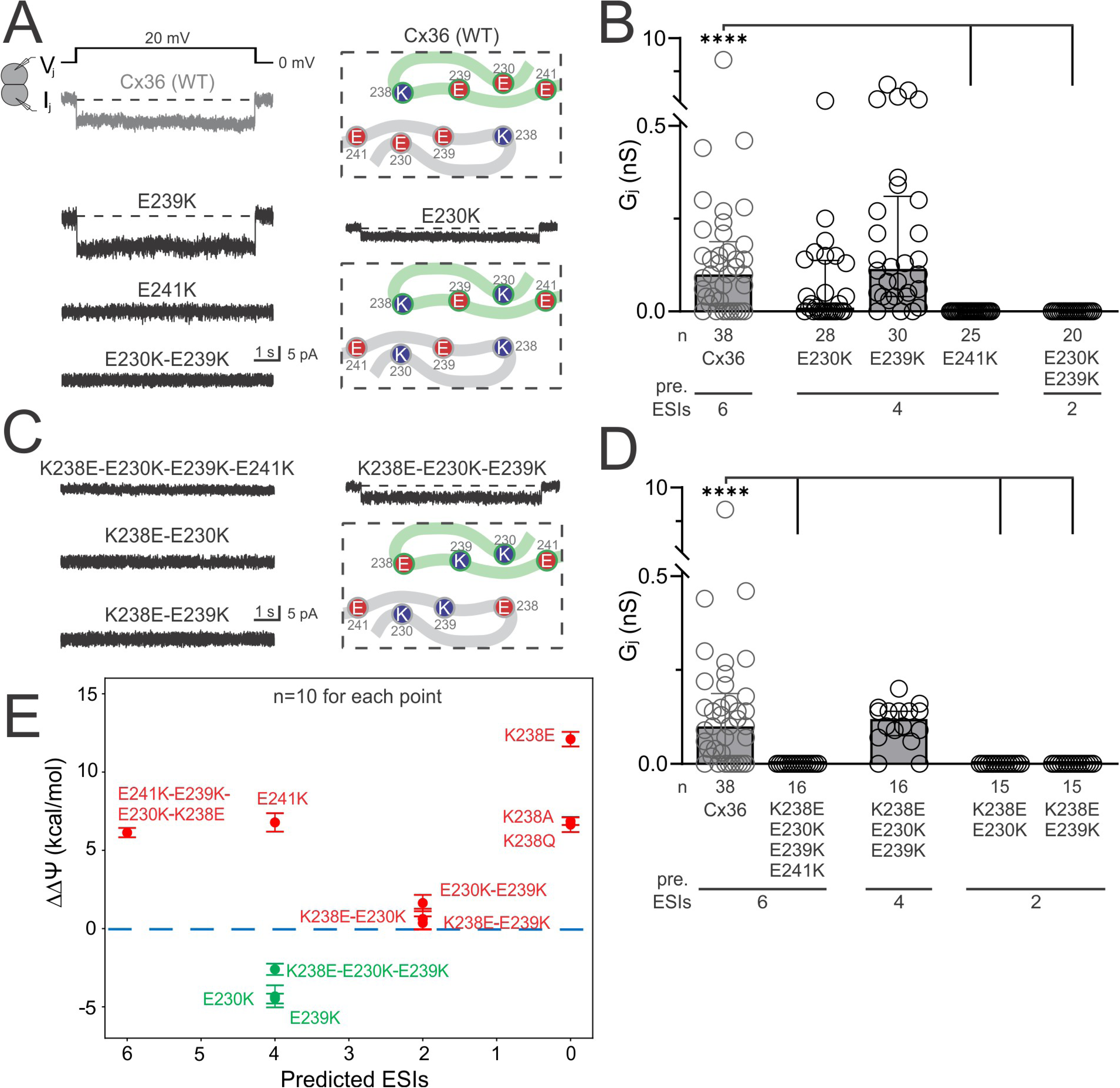
Functional status of Cx36 variants designed to reduce or reverse E2-E2 ESIs. (A, C) Representative I_j_s were recorded in response to a V_j_ pulse applied to DKO HEK293 cell pairs expressing a Cx36 variant designed to reduce (A) or reverse (C) the ESIs. Schematic drawings for the docking interface are shown for wildtype Cx36, E230K, E241K, K238E-E230K-E239K and K238E-E230K. For the predicted number of ESIs of these variants, see Fig. S2. (B, D) Bar graph summarizing the median G_j_ (error bars represent IQR) recorded for all cell pairs expressing variants designed to reduce (B) or reverse (D) the ESIs. The number of predicted ESIs per E2-E2 is indicated for each group below the X-axis (pre. ESIs). Statistical significance (\*\*\*\**P* < 0.0001) is obtained from Kruskal-Wallis tests followed by Dunn’s multiple comparisons tests comparing the G_j_ of each variant to that of Cx36. “n” is the number of biologically independent cell pairs for each group. I_j_ and G_j_s for Cx36 are shown here (same as Fig. 1) in grey for comparison purposes. (E) *In silico* calculations show that the change in electrostatic energy (ΔΔΨ) effectively distinguishes functional (green) from non-functional gap junction variants (red). Error bars indicate the standard deviation of the 10 independent runs of each variant.

Next, we reversed the charges of these residues to test whether the Cx36 variants could form functional GJs. Variants were designed and tested for their GJ function via dual patch clamp, including a fully reversed quadruple variant, K238E-E230K-E239K-E241K, and three partially reversed variants: K238E-E230K-E239K, K238E-E230K, and K238E-E239K (Fig. 2 and see Fig. S2). I_j_s recorded in response to a V_j_ pulse were only observed in cell pairs expressing K238E-E230K-E239K, but not in other variant-expressing cell pairs (Fig. 2C). The median G_j_ (IQR) of all cell pairs recorded was summarized in a bar graph (Fig. 2D, Table 2). The G_j_ of K238E-E230K-E239K was not significantly different from that of Cx36, while K238E-E230K-E239K-E241K, K238E-E230K, and K238E-E239K displayed no coupling (***P < 0.001 compared to Cx36), indicating that a partial reversal of ESIs with the K238E-E230K-E239K variant allowed for functional GJ formation.

Cx36 variants designed to eliminate (Fig. 1), reduce (Fig. 2A, B), or reverse (Fig. 2C, D) E2-E2 ESIs were plotted according to their predicted number of ESIs and the change in electrostatic interaction energy relative to the wildtype Cx36 GJ (ΔΔΨ, Fig. 2E). It is interesting to note that all variant GJs with ΔΔΨ above zero (i.e. increased electrostatic interaction energy relative to wildtype) were unable to form functional GJs in our patch clamp experiments, while all three variant GJs with ΔΔΨ below zero (i.e. decreased electrostatic interaction energy relative to wildtype) successfully formed functional GJs in patch clamp experiments (Fig. 2E). Overall, it appears that ΔΔΨ is an excellent predictor for the ability of any given Cx36 variant to form functional GJs. Although the number of ESIs was not a perfect predictor for functionality, all the variants with two or fewer predicted ESIs were unable to form any functional homotypic GJs, while the variants with four predicted ESIs were able to form functional GJs, except E241K.

### E241K-containing variants could not form functional GJs

The quadruple (K238E-E230K-E239K-E241K) and the single E241K Cx36 variants were predicted to have 6 and 4 estimated number of ESIs per E2-E2, respectively, yet both could not form functional GJs (Fig. 2). We further evaluated the GJ function of other triple or single variant combinations containing E241K in homo- or hetero-typic GJs, as shown in Fig. 3. All of these E241K-containing variants were also unable to form functional GJs (Fig. 3A, B, Table 2). A closer inspection of the Cx36 structure model revealed that, unlike E230, E239, and K238, which are at the surface of the GJ channel wall, E241 is buried in the channel wall structure surrounded by other nearby residues (Fig. 3C). E241K not only changed the charge of the sidechain from negative to positive, but also increased the size of the sidechain, which could cause steric clashes leading to structural changes at the E2 docking interface to impair docking. This explanation was supported by the fact that the missense variant E241A with a much-reduced sidechain size was found to successfully form functional GJs similar to that of Cx36 (Fig. 3A, B). We also studied whether the Cx36 E241K variant was able to form morphological GJ plaques using fluorescent protein-tagged variants, i.e. E241K-GFP and E241K-RFP. Unlike those found in Cx36-GFP and Cx36-RFP expressing cell pairs, E241K-GFP and E241K-RFP cell pairs rarely formed yellow morphological GJ plaques (Fig. 3D, E). This result is consistent with the idea that E241K and E241K-containing variants might alter the E2 docking structure, resulting in an inability to form functional GJs. Due to this possible docking structure change with E241K, this variant and any variants containing E241K were not included in the subsequent experiments and final data summary.

**Figure 3.**
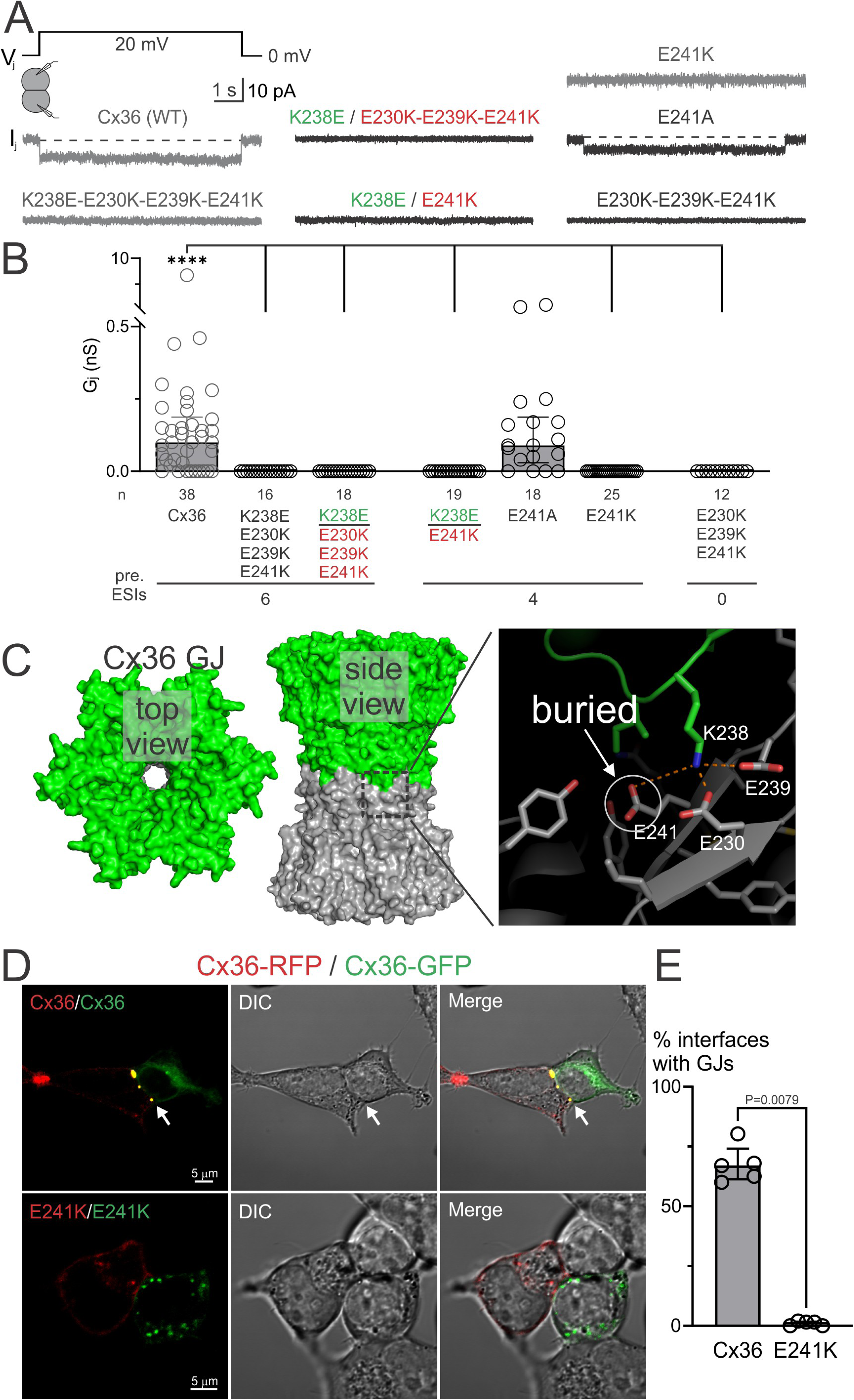
E241K-containing variants did not form functional GJs or morphological GJ plaques. (A) Representative I_j_s were recorded in response to a V_j_ pulse applied to DKO HEK293 cell pairs expressing a Cx36 variant(s) as indicated. (B) Bar graph summarizing the median G_j_ (error bars represent IQR) for all cell pairs recorded. The number of predicted ESIs per E2-E2 is indicated for each group below the X-axis (pre. ESIs). Statistical significance (\*\*\*\**P* < 0.0001) is obtained from a Kruskal-Wallis test followed by Dunn’s multiple comparisons tests comparing the G_j_ of each variant to that of Cx36. “n” is the number of biologically independent cell pairs for each group. I_j_s and G_j_s for Cx36, E241K, and K238E-E230K-E239K-E241K are shown here (same as Fig. 1 and 2) in grey for comparison purposes. (C) Cx36 GJ structure model (PDB:8XGD) in surface top view and side view to show the channel surface and wall (left and middle panels). The right panel shows a zoom-in view of a few residues near the docking interface. K238, E230, and E239 are located at the channel wall surface, with E241 buried in the channel wall protein (surrounded by a few other nearby residues). (D) Fluorescent, DIC, and merged images showing the localization of Cx36-RFP and Cx36-GFP, or E241K-RFP and E241K-GFP cell pairs in DKO HEK293 cells. Yellow GJ plaques were frequently observed at the cell-cell interface in cell pairs expressing Cx36-RFP and Cx36-GFP, but not in those expressing E241K-RFP and E241K-GFP. (E) Bar graph summarizing the median (error bars represent IQR) percentage of cell-cell interfaces that formed morphological GJ plaques for Cx36 and E241K cell pairs. Each data point (n=5) represents an independent transfection, where the percentage of cell-cell interfaces that formed morphological GJ plaques was counted from a minimum of 50 cell-cell interfaces for each group. A Mann-Whitney test was used to compare the percentage of cell-to-cell interfaces with morphological GJ plaques in Cx36 cell pairs, to GJ plaques formed in E241K cell pairs (*P*=0.0079).

### ESI number at the E2-E2 interface governs GJ formation

To investigate the minimum required ESIs to form functional Cx36 GJs, we used dual patch clamp to evaluate the function of additional Cx36 variants in heterotypic combinations (each cell expresses a different variant), ranging from five-to-one predicted ESIs per E2-E2 (Fig. S3, Table 2). All Cx36 variant combinations (homo- and heterotypic GJs) we tested were summarized in a table organized according to their ability to form functional GJs and the number of predicted ESIs per E2-E2 (Fig. 4A), with E241K-containing variant combinations excluded. A bar graph showing the median G_j_ (IQR) for cell pairs recorded is shown in Fig. 4B. Cx36 variant combinations predicted to form four or more ESIs per E2-E2 showed G_j_s that were similar to those of Cx36, demonstrating that they formed functional GJs. In comparison, variant combinations predicted to form two or fewer ESIs per E2-E2 showed G_j_s significantly lower than that of Cx36, indicating they were unable to form functional GJs. Among variant combinations predicted to form three ESIs, Cx36/K238A, Cx36/K238Q, E239K/E230-E239K, and Cx36/K238E-E239K expressing cell pairs demonstrated GJ function, while E241A/E230K-E239K and E230K/E230K-E239K did not readily show GJ coupling. Overall, our dual patch clamp data across 25 Cx36 variant GJ combinations suggests that a minimum of three predicted ESIs per E2-E2 are required but are not necessarily sufficient for functional GJ formation.

**Figure 4.**
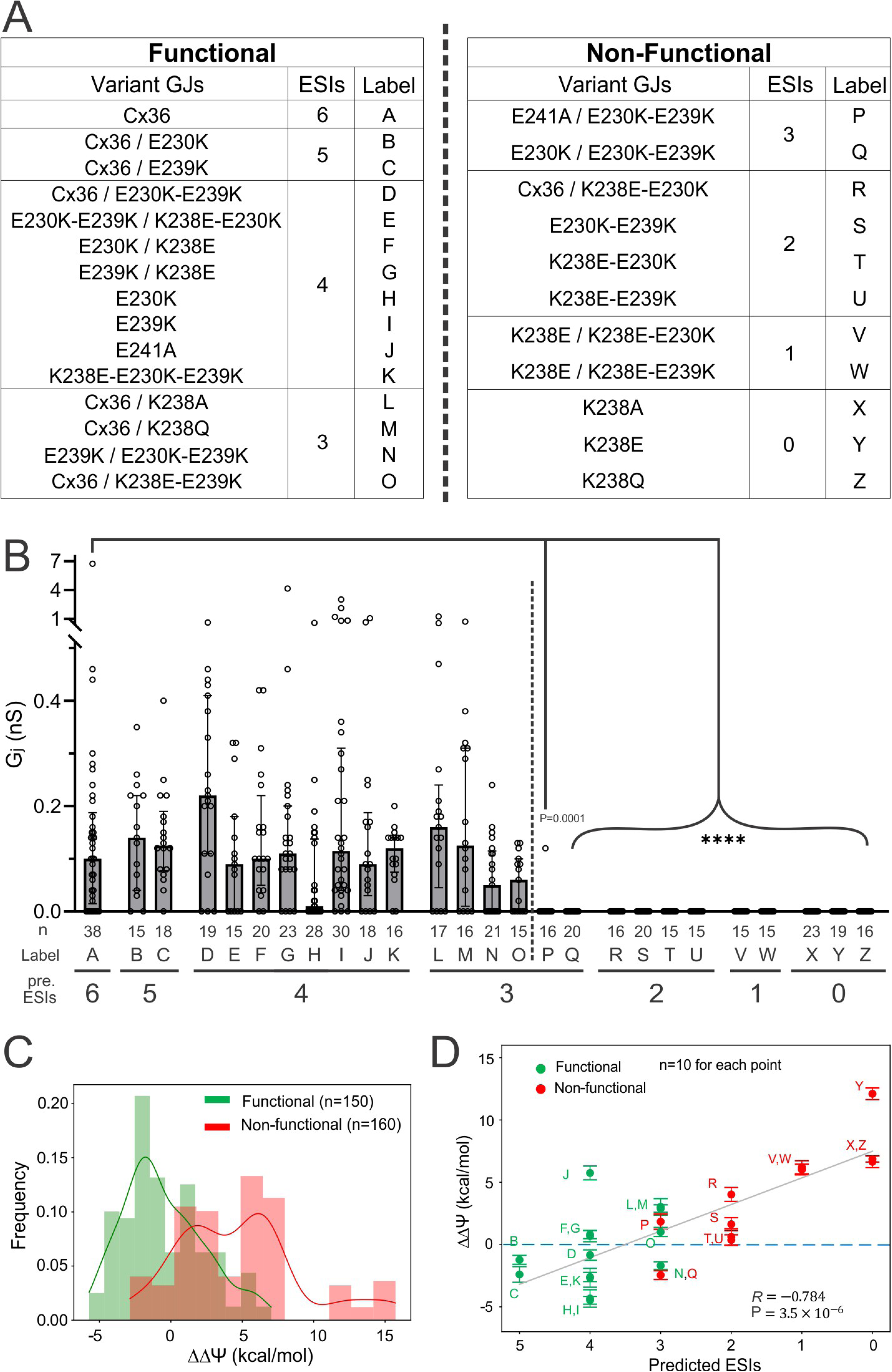
Functional status of Cx36 variant homo- and hetero-typic combinations. (A) All the homo- and hetero-typic Cx36 variant combinations except E241K and E241K-containing variants are organized according to functional outcome and ranked by predicted number of ESIs per E2-E2. Each variant combination is labeled with a capital letter linking to its median Gj as shown in (B). (B) Bar graph summarizing the median G_j_ (IQR) for all tested combinations. The number of measured biologically independent cell pairs (n) and the number of predicted ESIs per E2-E2 (pre. ESIs) are marked along the X-axis. Statistical significance (\*\*\*\**P* < 0.0001) is obtained from a Kruskal-Wallis test followed by Dunn’s multiple comparisons tests comparing the G_j_ of each variant to that of Cx36. (C) The distributions of ΔΔΨ of Cx36 variants show that the functional status of Cx36 variants can be primarily distinguished by ΔΔΨ — the changes in electrostatic energetics induced by the mutation. (D) Variants in (A) show a linear correlation between their predicted ESI and *in silico* calculated ΔΔΨ, with a Pearson correlation of -0.784. Error bars represent the standard deviation among 10 different runs. Note that X-axis is plotted based on decreasing number of ESIs.

We then ranked all the variant GJs by their electrostatic energy changes (i.e. ΔΔΨ, Fig. 4C). The functional variant GJs displayed a lower ΔΔΨ (green histogram and fitting curve), while non-functional variant GJs showed a higher ΔΔΨ (red histogram and fitting curve, Fig. 4C). No other contributing energy term by comparison showed greater ability to distinguish the variant functional status than the electrostatic energy (Fig. S4A), further supporting the importance of electrostatic interactions at the E2-E2 in Cx36 GJ function. Plotting the predicted ESIs with the ΔΔΨ (Fig. 4D) demonstrated that these two parameters are correlated with a Pearson correlation coefficient of -0.784, indicating that our predicted number of ESIs is a valid estimation of the electrostatic energy change (ΔΔΨ). Furthermore, the linear fitting line crosses zero ΔΔΨ (i.e. change in electrostatic energy changes from lower than wildtype to higher than wildtype Cx36) between three and four predicted ESIs, aligning nicely with our functional data where variant GJs predicted to form four or more ESIs formed functional GJs, variant GJs predicted to form two or fewer ESIs did not form functional GJs, and only some variant GJs predicted to form three ESIs formed functional GJs, while others did not. Overall predicted number of ESIs together with ΔΔΨ calculations proves to be a good predictor of Cx36 variant function. A similar plot of predicted ESIs but with a change in Gibbs free energy (ΔΔG) instead of a change in electrostatic energy for all variant GJs demonstrated a similar linear correlation (Pearson correlation coefficient of -0.768; Fig. S4B) to that shown in Fig. 4D, supporting the hypothesis that altering the charged docking residues may effectively alter the variant GJ binding energy and function.

To remove potential bias arising from the specific crystal structure used, we conducted analogous analyses on another experimentally derived E2-E2 complex (PDB ID: 8IYG). Similar significant correlations between ΔΔG, ΔΔΨ, and the number of ESIs were observed (Pearson correlation coefficient: -0.734 and -0.781, respectively; Fig. S5A, B), reinforcing that electrostatic interactions predominantly drive binding energetics and are critical determinants of functional integrity in the E2-E2 protein complex (Fig. S5C).

### Human Cx36 exhibits minimal heterotypic GJ function with other brain connexins

To evaluate whether human Cx36 is capable of forming functional heterotypic GJs with other connexins, we paired Cx36 expressing cells (DsRed positive) with cells expressing one of the following connexins: Cx26, Cx30, Cx31.3, Cx32, Cx43, Cx45, or Cx47 (GFP positive). GJ coupling between cell pairs was assessed using dual whole-cell patch clamp recordings. Representative junctional currents (I_j_s) in response to a V_j_ pulse are shown in Fig. 5A. In nearly all heterotypic combinations tested, no measurable junctional currents were detected. The median junctional conductance (G_j_) values, along with their interquartile ranges (IQRs), are summarized in Fig. 5B and Table 3. Across all combinations, G_j_ values for heterotypic pairs were significantly lower than those measured for homotypic Cx36 GJs (***P < 0.001), confirming that Cx36 is largely incompatible with other major brain connexins for functional heterotypic GJ formation under our experimental conditions. As an alternative approach, we artificially constructed a structural model by substituting one hemichannel in the resolved Cx36/Cx36 GJ structure with a Cx32 hemichannel. After a structural relaxation in molecular dynamics simulations, we calculated the electrostatic energy across the E2–E2 docking interface. This analysis revealed that the electrostatic interaction energy for the Cx36/Cx32 GJ interface is approximately −0.3 kcal/mol, compared with −3.0 kcal/mol for the Cx36/Cx36 GJ interface, while the Cx32/Cx32 GJ interface exhibits a near-neutral electrostatic contribution (∼0.1 kcal/mol). These results indicate that the electrostatic interaction network that stabilizes Cx36 docking is substantially reduced in the artificial model of Cx36/Cx32 GJ.

**Figure 5.**
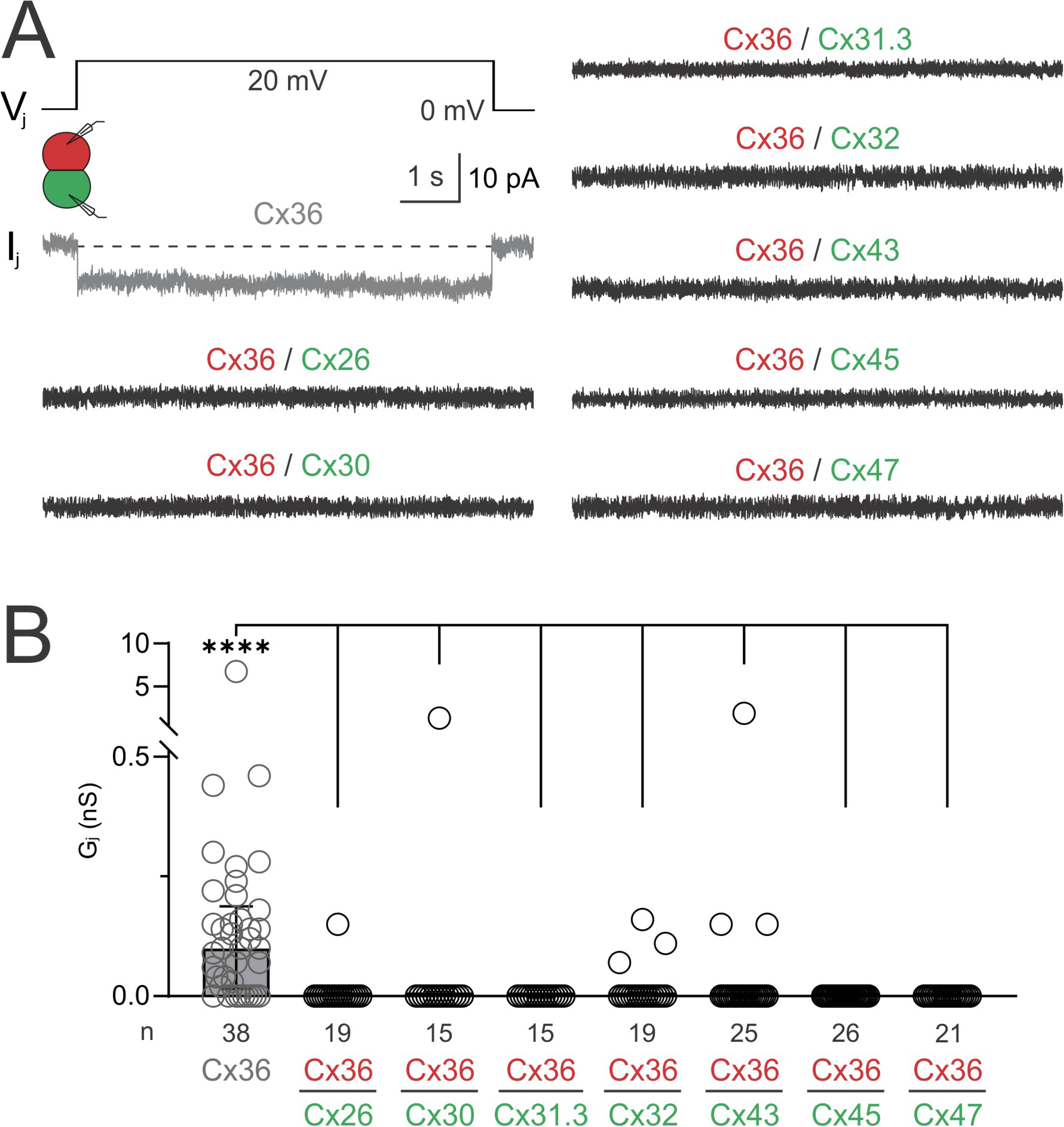
Cx36 exhibits minimal functional heterotypic GJ function with other brain connexins. (A) Representative I_j_s were recorded in response to a V_j_ pulse applied to DKO HEK293 cell pairs expressing Cx36 in one cell (DsRed positive), and Cx26, Cx30, Cx31.3, Cx32, Cx43, Cx45, or Cx47 in the other cell (GFP positive) as indicated. (B) Bar graph summarizing the median G_j_ (error bars representing IQR) for all cell pairs recorded. A Kruskal-Wallis test followed by Dunn’s multiple comparisons tests were performed to compare the G_j_ of each heterotypic GJ to that of Cx36 where in all cases, heterotypic G_j_s were significantly lower (****P < 0.0001). “n” represents the number of biologically independent cell pairs measured for each group. I_j_ and G_j_s for Cx36 are shown here (same as Fig. 1) in grey for comparison purposes.

**Table 3.** Median G_j_s (IQR) for Cx36 heterotypic combinations with other brain connexins, and their respective homotypic GJs.

| **G<sub>j</sub>s** | **Median G<sub>j</sub> (nS)** | **IQR** | **# cell pairs** | **# trans.** |
| --- | --- | --- | --- | --- |
| Cx36 / Cx26 | 0 | 0 - 0 | 19 | 3 |
| Cx36 / Cx30 | 0 | 0 - 0 | 15 | 3 |
| Cx36 / Cx31.3 | 0 | 0 - 0 | 15 | 4 |
| Cx36 / Cx32 | 0 | 0 - 0 | 19 | 5 |
| Cx36 / Cx43 | 0 | 0 - 0 | 25 | 6 |
| Cx36 / Cx45 | 0 | 0 - 0 | 26 | 6 |
| Cx36 / Cx47 | 0 | 0 - 0 | 21 | 5 |
| Cx26 / Cx26 | 1.96 | 0.04 - 4.47 | 16 | 5 |
| Cx30 / Cx30 | 6.55 | 0.43 - 21.28 | 16 | 4 |
| Cx31.3 / Cx31.3 | 0 | 0 - 0 | 17 | 5 |
| Cx32 / Cx32 | 0.85 | 0 - 16.23 | 17 | 5 |
| Cx43 / Cx43 | 0.46 | 0 - 9.93 | 17 | 3 |
| Cx45 / Cx45 | 6.80 | 0.5 - 19.6 | 29 | 10 |
| Cx47 / Cx47 | 1.30 | 0.08 - 7.68 | 20 | 6 |

“IQR”, interquartile range; “# cell pairs” represents the number of cell pairs recorded for each heterotypic G<sub>j</sub>; “# trans.” represents the number of transfections.

To ensure these connexins are capable of assembling into hemichannels and trafficking to the plasma membrane, we tested the ability of these connexins to form functional homotypic GJs. Each of these connexins, with the exception of Cx31.3, successfully formed functional GJs with expected V_j_-gating signatures of the respective connexin GJs (Fig. S6), demonstrating that these connexins readily formed hemichannels on the cell plasma membrane, a prerequisite for forming functional GJs. The inability of Cx31.3 to form functional GJs in our model cells is expected, as similar results were obtained using different model cells^43^. We also studied the localization of carboxyl-terminal GFP-tagged Cx31.3 (Cx31.3-GFP) in DKO HEK293 cells, which appeared to cluster in the cytosol and on the plasma membrane, similar to those of Cx43-GFP and Cx45-GFP, but different from that of GFP alone, which was localized evenly throughout the cytosol and nuclei (Fig. S7). The localization of Cx31.3-GFP is virtually the same as that described using anti-Cx31.3 antibody labeling in an earlier study, where Cx31.3 was found to be localized to the plasma membrane and formed functional hemichannels^43^.

## Discussion

Recent structure models of the human Cx36 (hCx36) GJ revealed novel electrostatic interactions (ESIs) between charged residues at the E2-E2 docking interface, with up to 6 predicted ESIs at each pair of docked E2-E2 interface or 36 ESIs at two docked hemichannels^39, 40, 41, 44^ (Fig. 1A). We systematically designed a series of variants on these putative ESI-forming residues, aiming to eliminate, reduce, or reverse these ESIs at the E2-E2 docking interface of Cx36 GJ. Computational calculations on the designed variant GJs showed that the change in electrostatic interaction energy (ΔΔΨ) positively correlated with the change in hemichannel binding free energy (ΔΔG). To test the roles of ESIs in an experimental setting, we used the dual patch clamp method to assess the ability of these Cx36 variants to form functional GJs in a connexin-deficient human cell line (genetically engineered HEK293)^42^. Our study, for the first time to our knowledge, demonstrated the importance of ESIs at the Cx36 E2-E2 docking interface for the formation of functional GJs, where a minimum of 3 predicted number of ESIs per E2-E2 (or 18 ESIs between 2 hemichannels) was required for these Cx36 variants to form functional GJs. Calculating the change in both binding free energy (ΔΔG) and electrostatic free energy (ΔΔΨ) associated with our designed E2 variants in different GJs showed a significant correlation with the predicted number of ESIs, indicating the number of ESIs is a valid predictor for the successful formation of functional GJs. The unique ESIs at the Cx36 E2 docking interface are also likely to play a role in its docking selectivity, as Cx36 rarely formed functional heterotypic GJs with any other brain expressed connexins, including Cx26, Cx30, Cx31.3, Cx32, Cx43, Cx45, and Cx47. A comprehensive understanding of the mechanisms of neuronal Cx36 docking selectivity is very important for its unique, highly restricted interneuronal communication with little cross-talking via heterotypic GJs to other cell types in the brain.

Several of our designed Cx36 variants in either homotypic or heterotypic GJs with 3 or fewer predicted ESIs at each of the six E2-E2 docking interfaces were unable to form functional GJs in cell pairs expressing these variants. One possibility of this failure might not be due to insufficient ESIs to allow two hemichannels to dock together, but might be attributed to a defective trafficking of these variant hemichannels, so that they’re unable to reach the plasma membrane, and therefore not available at the cell surface for docking with hemichannels on neighbouring cells. A closer look at our experimental results for these variants, which were unable to form functional homo- or heterotypic GJs (Fig. S8), indicates that they were all able to successfully form functional GJs in at least one other heterotypic GJ where a higher number of E2-E2 ESIs was predicted (see Fig. S8). The successful formation of functional GJs demonstrated that these variant hemichannels were able to traffic to the plasma membrane and were functionally competent when an appropriate docking partner was available. Thus, the lack of GJ formation in these variant GJs is more plausibly explained by the insufficient electrostatic interactions (with lower predicted ESIs at the E2-E2 docking interface), which are critical for stabilizing their hemichannel docking and formation of functional GJs.

In the past 2 years, four Cx36 GJ structures have been resolved at high resolution using a Cryo-EM approach from three independent research groups^39, 40, 41, 44^. All of these Cx36 GJ structure models showed similar structures at the docking interface between the extracellular domains (i.e. E1-E1 and E2-E2). Though all these GJ structures proposed to have electrostatic interactions (ESIs) at the E2-E2 docking interface, the number of ESIs for each pair of docked E2-E2 varied from 2 to 6, depending on the distances of K238 to the docked triple glutamate residues (E230, E239, E241) in these static GJ models^39, 40^. However, we found that with some minor changes in the orientation of the involved residue sidechains, the sidechain of K238 is close enough (within 4 Å) to make ESIs with each of the three glutamate residues in every Cx36 GJ structure. Considering the dynamic nature of native proteins, especially on these long sidechain residues (lysine and glutamate), we believe that K238 on one E2 is likely to interact with each of the 3 glutamate residues at the docked E2, forming up to 3 ESIs (or 6 ESIs total) for each E2-E2 docking interface (due to rotational symmetry, see Fig. 1A).

Previous studies on purified Cx36 oligomers showed that complete elimination of ESIs at the E2-E2 docking interface with mutations (such as K238E or K238A) substantially decreased the ratio of dodecamers to hexamers based on size exclusion chromatography, suggesting an impairment in the docking of Cx36 hemichannels^39, 40^. Reduction of ESIs by mutating each of the glutamate residues into positively charged or neutral residues (e.g. E230K or E230A) showed a moderate decrease in the ratio of dodecamer / hexamer, indicating a partially impaired Cx36 docking^39^. Here, we generated a series of variants on these key docking residues in Cx36 and assessed the functional consequences of modifying different numbers of ESIs in Cx36 GJ in homotypic and heterotypic GJs. Consistent with previous findings on purified protein oligomers, the total elimination of ESIs with K238E/A/Q prevented the formation of any functional GJs. Reduction of ESIs from predicted 6 to 4 with designed Cx36 variants had no detectable change in the formation of functional Cx36 GJs (except for E241K and variants containing this missense mutation), while further reduction of ESIs to 2 or 1 resulted in variants unable to form any functional GJs (Fig. 4). We were very excited that our designed variant with reversed ESIs (K238E-E230K-E239K) at the E2 docking interface were able to form functional GJs, demonstrating the importance of ESIs for Cx36 GJ docking (Fig. 2). Through systematic investigation of more than 2 dozen different combinations of GJs with various predicted ESIs, we found that 3 ESIs appear to be the minimum number of ESIs to form functional GJs. It is interesting to note that a recent paper reported engineering a pair of variants in fish Cx36 homologues to be heterotypic docking compatible, but not able to form homotypic GJs and also unable to form heterotypic GJs with Cx36 and Cx43, consistent with our findings but using very different approaches and *in vitro* or *in vivo* models^45^.

Among our tested 6 heterotypic GJs with 3 predicted ESIs, 4 of them formed functional GJs and 2 of them could not, indicating that 3 ESIs at the E2-E2 docking interface are the minimum requirement, but not sufficient for the formation of functional GJs. The underlying structural mechanisms are not fully clear, but we believe that co-operative ESIs could be formed at the docking interface for some of these GJs, promoting the formation of functional GJs. Multiple ESIs formed between the positively charged lysine and 3 glutamate residues at the E2 docking interface of Cx36 are likely cooperative^46^, as the angles between Cα atoms are less than 90°, which is true for the Cx36 variant GJs. The net strength of co-operative ESIs is more than the sum of the energies of individual pair or triple ESIs. We believe that co-operative ESIs might play a role in the formation of functional GJs in some of the GJs with 3 predicted ESIs (for example, Cx36 / K238A and Cx36 / K238Q), but additional factors could also contribute to the reason why 2 heterotypic GJs could not form functional GJs even with 3 predicted ESIs (E241A / E230K-E239K; E230K / E230K-E239K). One possibility is that the three glutamate residues on the E2 do not serve equal roles in anchoring two docked E2 domains. Perhaps E239 plays a slightly less important role relative to E230 and E241, as the heterotypic E239K / E230K-E239K GJ formed functional GJs, while E241A or E230K docked with E230K-E239K were unable to form functional GJs. This idea is also consistent with the observations that 1) in skate Cx35, an ortholog of Cx36, a methionine (M219) is at the equivalent position of E239, not a negatively charged residue; 2) in the phylogenomic database Orthologous MAtrix (OMA) for Cx36, there are 154 sequences from different species and their E2 sequence logo showed that the position of E239 is the only one among the three glutamate residues that showed some variation with ∼2/3 species with glutamate (E) and ∼1/3 species with aspartate (D), indicating that this position could tolerate variation in its sidechain length. Additional experiments are needed to clearly demonstrate if there are different roles for the triple glutamate residues on the E2 of Cx36.

Using FoldX 5.0^59^, we analyzed all the tested variant GJs for their change in Gibbs free energy (ΔΔG) from the wild-type Cx36 GJ and revealed a linear correlation with the number of ESIs. Further analyzing each of the composed energy components, we found that the changes in electrostatic energy (ΔΔΨ) were the most dominant energy term for the correlation ΔΔG and, as predicted, also showed significant correlation to the number of ESIs, indicating ESI number is a reliable parameter for estimating Cx36 variant structure stability and likelihood of formation of functional GJs. Our findings revealed similar correlations for two independent Cx36 structure models (PDB IDs: 8XGD and 8IYG, as shown in Fig. 4, S4, and S5), indicating that the calculations are broadly applicable and independent of the initial structure models.

Cx36 is widely expressed in the brain, forming GJs (also known as electrical synapses) to synchronize neuronal network activities and rhythmic oscillations, while its expression in pancreatic β islets synchronizes cell activity to regulate insulin secretion^16, 17, 47^. The importance of Cx36 GJs has been demonstrated in mouse models with the Cx36 gene (*GJD2*) ablated, which showed a variety of impairments, including in learning and memory, reduced synchronized inhibitory neuronal activities and an elevated risk of sudden death during the neonatal stage^11, 48, 49, 50^. *GJD2* has also been linked to epilepsy, inflammation, amyotrophic lateral sclerosis, and diabetes^51, 52, 53, 54^. Multiple disease-linkage of *GJD2* is consistent with findings from a large cohort of human genome sequencing results (gnomAD v4.1.0), where less than half (12/24.4) of the predicted loss of function mutants in *GJD2* were observed in the database. Similarly, only 63% (268/424.8) of the expected missense variants were observed, indicating this gene may be under negative selection, and is likely intolerant to loss of function and missense variants, which would explain why they are less frequently observed in the database than expected^55^. In contrast, the number of synonymous variants observed was very close to the expected number of variants (https://gnomad.broadinstitute.org/gene/ENSG00000159248?dataset=gnomad_r4).

Our results for human Cx36 docking compatibility are similar to those found in rodent Cx36, where Cx36 can dock with Cx36 to form functional homotypic GJs, but is unlikely to dock with other connexins to form heterotypic GJs^21, 22^. We tested the ability of Cx36 to dock with connexins expressed in other neurons (Cx45), astrocytes (Cx26, Cx30, Cx43), oligodendrocytes (Cx31.3, Cx32, Cx47), and microglia (Cx26, Cx32, Cx43)^5^, which supports the idea of unique selective intercellular communication mediated by Cx36 GJs (Fig. 5). Perhaps nature designed Cx36 to be more restrictive in terms of docking compatibility to ensure specific passage for inter-neuronal communication with virtually no GJ linkage to other brain cells expressing other connexins. Docking compatibility for connexins is very critical in establishing intricate networks of GJs mediating selected signals and, at the same time, preventing unnecessary crosstalk to ensure normal physiology.

In conclusion, Cx36 GJs exhibit ESIs between charged residues at the docked E2-E2 interfaces, distinct from the hydrogen bond interactions present at the E2-E2 interfaces of all other human connexins. Here, we quantitatively evaluated the contribution of these ESIs to docking and binding energetics in functional Cx36 GJs. Our results reveal that the minimum number of ESIs in Cx36 GJs required to form functional GJs is 3 / E2-E2 or 18 / GJ in Cx36 GJs. The unique ESIs in Cx36 GJ are critical for Cx36 to form functional GJs with itself and may also play a role in specifically targeted GJ communication between neurons in the brain and pancreatic β cells in the pancreas, with virtually no cross communication to other connexins expressed in the same tissue.

## Materials and Methods

### Plasmid construction

The cDNA for human Cx36 was subcloned into an untagged pIRES2-DsRed-Express2 expression vector between the restriction enzyme sites Nhel, and EcoRI. Similarly, the cDNA for human Cx30, Cx31.3, Cx43, and Cx47 were subcloned into untagged pIRES2-EGFP expression vectors between the restriction enzyme sites Nhel and BamHI (Cx30), Xhol and BamHI (Cx31.3), EcoRI and BamHI (Cx43), and Xhol and BamHI (Cx47). pIRES2-EGFP-hCx36, pIRES2-EGFP-hCx26, and pIRES2-EGFP-hCx45 expression vectors were generated as described previously^37, 42,56^. For morphological GJ plaque formation experiments, human Cx36 was subcloned into an expression vector pEGFP-N1 (or pTaqRFP-N) with a fluorescent protein tag in frame linked at the carboxyl terminus of Cx36, between restriction sites Nhel, and EcoRI. The stop codon of Cx36 was removed for both Cx36-GFP and Cx36-RFP. A peptide linker (with 29 residues, KGAPVYSRVDTRGILQSTVPRARDPPVAT) was in between Cx36 and GFP (the same linker was in between Cx36 and RFP). For the localization of Cx31.3, Cx31.3 was subcloned into a pEGFP-N1 expression vector with a fluorescent GFP tag in frame linked at the carboxyl terminus of Cx31.3 (Cx31.3-GFP), between restriction sites Xhol and BamHI. The stop codon of Cx31.3 was removed, and a peptide linker (WDPPVAT) was observed between Cx31.3 and GFP. For the localization of Cx43 and Cx45, we used Cx43-GFP and Cx45-GFP, which were generated as described previously^56, 57^.

pIRES2-EGFP-hCx36, pIRES2-DsRed-Express2-hCx36, pEGFP-N1-hCx36, or pTaqRFP-N-hCx36 were used as templates to generate the following single, double, triple and quadruple missense Cx36 variants using the primers as listed in Table 1. Note that some of the double, triple, or quadruple mutants were generated on single, double, or triple variants as indicated.

### Cell culture and transient transfection

Connexin-deficient Human Embryonic Kidney (HEK293-AD) cells were purchased from Agilent (catalog part# 240085) and genetically engineered to ablate Cx43 and Cx45 using CRISPR-cas9^42, 58^. The resulting double knockout (DKO) HEK293 cells were used to study the function of gap junctions formed by recombinantly expressed human connexins. DKO HEK293 cells were grown in Dulbecco’s modified Eagle’s medium (DMEM) (Life Technologies, Cat. No. 10313-021) containing 4.5 g/L D-glucose, and 110 mg/L sodium pyruvate, and supplemented with 10% fetal bovine serum (Life Technologies, Cat. No. 080150), 1% penicillin streptomycin (Life Technologies, Cat. No. 15140-122), and 1% GlutaMAX (FBS, Life Technologies, Cat. No. 35050-061) in an incubator with 5% CO2 at 37°C.

DKO HEK293 cells were transfected at a 1:2 ratio of cDNA construct to X-tremeGENE HP DNA transfection reagent (Roche Diagnostics GmbH, Cat. No. 06366546001), ranging from 0.5-0.8 μg of cDNA construct and 1.0-1.6 μL of transfection reagent, in Opti-MEM + GlutaMAX-I medium (Cat. No. 51985-034) for 5 h in an incubator with 5% CO_2_ at 37°C. Following transfection, the growth medium was changed back to FBS-containing DMEM. Cells were then replated onto glass coverslips and incubated with 5% CO_2_ at 37°C for 3-20 h prior to patch clamp recording. A range of replating time points was used for different connexins and connexin variants to observe the optimal time for GJ formation in cell pairs. Isolated cell pairs positively transfected with a single connexin or connexin variant (with successful expression of a fluorescent reporter, GFP or DsRed) were selected for the study of homotypic GJs. For experiments with heterotypic cell pairs, we mixed cells transfected with a connexin variant (with GFP reporter), with another connexin variant (with DsRed reporter) and isolated cell pairs with one expressing GFP and the other expressing DsRed were selected for patch clamp recording. For morphological GJ plaque formation and connexin localization experiments, transfected cells were replated onto glass coverslips and incubated with 5% CO_2_ at 37°C for 5-18 h prior to visualization under a confocal microscope (see below).

### Electrophysiological recording

Glass coverslips with transfected cells were placed into the recording chamber of an upright microscope (Olympus BX51WI, Evident Corporation, Shinjuku, Tokyo, Japan), where positively transfected cell pairs were selected for dual whole cell voltage clamp to assess GJ function. The recording chamber was filled with extracellular solution (ECS) containing (in mM): 135 NaCl, 2 CsCl, 2 CaCl_2_, 1 MgCl_2_,1 BaCl_2_,10 HEPES,5 KCl, 5 D-(+)-glucose, 2 sodium pyruvate, pH adjusted to 7.4 with 1 M NaOH, and osmolarity of 310–320 mOsm/L. Patch pipettes pulled using a micropipette puller (PC-100, Narishige International, Amityville, NY) were filled with intracellular solution (ICS) containing (in mM): 130 CsCl, 10 EGTA, 0.5 CaCl_2_, 3 MgATP, 2 Na_2_ATP, 10 HEPES, adjusted to pH 7.2 with 1 M CsOH, and osmolarity of 290–300 mOsm/L. Dual whole cell voltage clamp was performed on cell pairs at room temperature (22 – 24°C) using a MultiClamp 700 A amplifier (Molecular Devices, Sunnyvale, CA). Each cell of the pair was initially voltage clamped at 0 mV, and voltage pulses were applied to one cell of the pair, ranging from ±20 to ±100 mV in steps of ±20 mV over 7 seconds to establish transjunctional voltage (V_j_). Gap junctional current (I_j_) was recorded in the other cell of the pair, which was continually held at 0 mV throughout each V_j_ step. The junctional current (I_j_) was low-pass filtered (Bessel filter at 1 kHz) and recorded using pClamp10.7 software at a sampling frequency of 10 kHz via an AD/DA converter (Digidata 1550, Molecular Devices, Sunnyvale, CA). Gap junctional conductance (G_j_) was calculated using the equation: 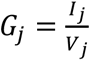

### Visualization of morphological GJ plaques and connexin localization

To assess morphological GJ plaque formation for Cx36 E241K, we used two different fluorescent protein-tagged E241K, i.e. E241K-GFP and E241K-RFP. Two different tagged E241K constructs were individually transfected into DKO HEK293 cells. After overnight incubation, these cells were mixed together and plated onto coverslips for 15-18 hours to allow formation of cell pairs with one expressing E241K-GFP and the other expressing E241K-RFP. Cx36-GFP and Cx36-RFP were used as positive controls. GFP and RFP vectors were used as negative control. Glass coverslips with transfected cells were observed under a confocal microscope (Zeiss LSM800 with Airyscan, Zeiss, Oberkochen, Germany) using a 40× water-immersion lens (NA 1.2) to assess for the formation of morphological GJ plaques commonly observed as bright yellow clusters at the cell-cell junction of GFP and RFP positive cells. The percentage of GFP and RFP positive cell-cell interfaces with bright yellow clusters (deemed to be morphological GJ plaques) was evaluated by counting at least 50 cell-cell interfaces for each transfection. Green and red fluorescent images were superimposed, along with a differential interference contrast (DIC) image for E241K and wild-type Cx36, similar to our earlier studies^35, 36^.

To assess the localization of Cx31.3, plasmid cDNA encoding carboxyl-terminal GFP-tagged Cx31.3 (Cx31.3-GFP) was transfected into DKO HEK293 cells. cDNAs of Cx43-GFP and Cx45-GFP were used as positive controls, and a GFP vector alone was used as a negative control. Glass coverslips with transfected cells were observed under a confocal microscope (Zeiss LSM800 with Airyscan) using a 40× water-immersion lens (NA 1.2) to assess the localization of each connexin-GFP, where representative green fluorescent images, along with DIC images were obtained.

### Free energy calculation on designed Cx36 variants

Point mutations were introduced into the wild-type Cx36 E2–E2 docking interface using FoldX (version 5.1)^59^. FoldX is an empirical force field originally developed to predict the effect of mutations on protein stability and interaction energies, utilizing a rigid-backbone approximation and detailed side-chain rotamer libraries. The protocol included mutating residues listed in Table 1 and their different combinations for homotypic and heterotypic GJs (Fig. S2), energy-minimizing the local environment, and computing the change in Gibbs free energy of folding (ΔΔG) relative to wild-type GJ. FoldX decomposes ΔG into key physical terms, including van der Waals contacts, solvation, hydrogen bonding, electrostatics, entropic costs, and so on.

To assess which energy terms best distinguish functional from non-functional Cx36 variant GJs, we computed the Silhouette score for each term extracted by FoldX. The Silhouette coefficient measures the cohesion and separation of clusters (here, functional vs. non-functional), defined for each data point i as:

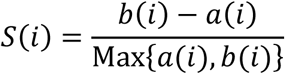

where a(i) is the mean intra-cluster distance and b(i) the mean nearest-cluster distance. Scores range from -1 (poor cluster assignment) to +1 (well-separated). We averaged scores for each energy term in both one-dimensional (single feature) and two-dimensional (feature pair) analyses. A higher average Silhouette score indicates a stronger ability of that term to differentiate functional status.

### Statistics and reproducibility

Gap junctional conductance (G_j_) data gathered using dual whole-cell patch-clamp is graphically represented as median G_j_ with error bars representing interquartile range [described as Median (interquartile range, IQR)]. Each independent cell pair measured is considered a biological replicate (n), where a minimum of 12 cell pairs were measured across at least 3 separate transfections for each experimental group. Cell pairs that had a G_j_ greater than 50 nS were excluded because it is impossible to differentiate the gap junction-coupled cell pairs from “apparently highly coupled cell pairs” due to a cytoplasmic bridge. Additionally, patch clamp recordings with input resistance under 200 MΩ were excluded from the G_j_ measurement due to possible poor seal quality associated voltage clamp errors. G_j_ data were analyzed using a two-tailed Kruskal–Wallis test followed by a Dunn’s post hoc test to compare data across multiple groups, or a two-tailed Mann-Whitney test to compare data between only two groups, as specified for each figure. Statistical significance is indicated on each bar graph (****P< 0.0001, or otherwise specified), and is focused on biologically meaningful group comparisons.

The percentage of positively transfected cell pairs with morphological GJ plaques is graphically represented as median (IQR). Each independent transfection (across 5 transfections for each experimental group) is considered an independent biological replicate, while at least 50 cell-cell interfaces were assessed for each experimental group per transfection. A two-tailed Mann-Whitney test was used to compare the percentage of morphological GJ plaque formation between Cx36-Cx36 and E241K-E241K cell interfaces. Statistical significance is indicated on the relevant bar graph.

## Acknowledgements

We thank Rania Amayem and Jennifer Zhu for technical help and discussions. We also thank Drs. Dale Laird and Peter Stathopulos for constructive discussions.

## Ethics Declarations

Competing Interests None to declare.

## Data Availability

Numerical source data underlying patch clamp and percentage of morphological GJ plaque bar graphs are provided in Supplementary Data 1. Raw patch clamp recordings are available upon reasonable request to the corresponding author. Computational data can be accessed through Zenodo at https://zenodo.org/records/20303481.

## Code Availability

All computational analysis scripts, datasets, and figure source files generated in this study are available at GitHub (https://github.com/zhaolabutmb/Cx36_Channel_Electrostatics), or Zenodo at (https://zenodo.org/records/20303481).

## Author Contribution

R.S.W. designed and performed most of the dual patch clamp experiments and localization experiments with tagged Cx36 variants, analyzed data, generated bar graphs, and wrote the first draft of the manuscript. Y.T.Z. designed and performed part of the dual patch clamp experiments and associated data analysis. H.C. designed and generated all cDNA constructs, performed Cx31.3 localization studies, and generated the associated figure. Z.S. calculated the free energies for the E2-E2 docking residues, wrote the computational session of the manuscript. H.Z. designed and supervised the computational calculations, wrote the computational session and critically revised the manuscript. D.B. designed the project, provided funding support, supervised the data analysis, and critically revised the manuscript.

