## supplementary_information for "Quantifying Electrostatic Control of Docking and Binding Energetics in Functional Cx36 Gap Junctions"

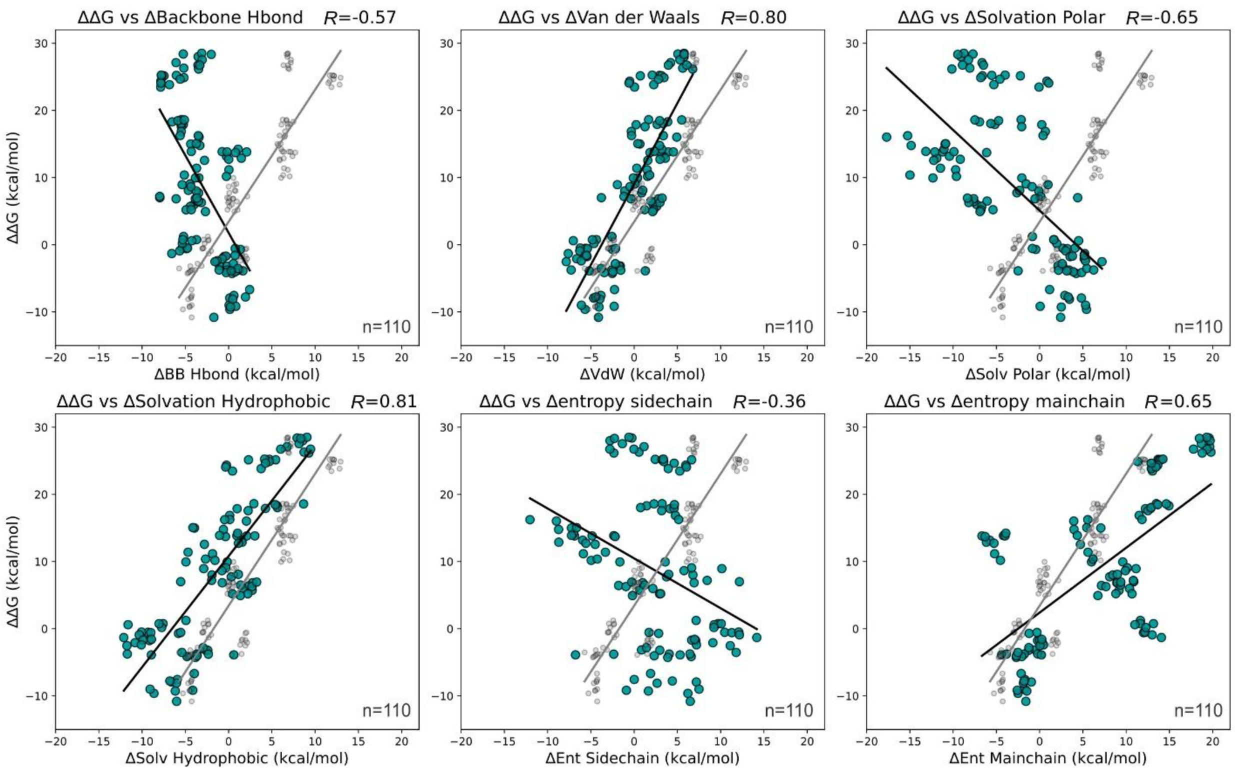


**Figure S1. Relationship between changes in binding free energy (ΔΔG) and individual contributing energy components**. Each of the six panels shows ΔΔG plotted against one of the six FoldX energy terms (panel titles indicate the specific component and its Pearson correlation coefficient). Each plot includes n = 110 samples obtained from 10 independent runs of energy calculation across 11 homotypic variants. Rp represents the Pearson correlation coefficient. These plots allow direct comparison of how strongly each energy component correlates with ΔΔG across the Cx36 mutants. Gray dots and the fitting line on each panel represent electrostatics (ΔΔΨ) plotted against ΔΔG for direct comparison purposes (replotted from Fig. 1B).

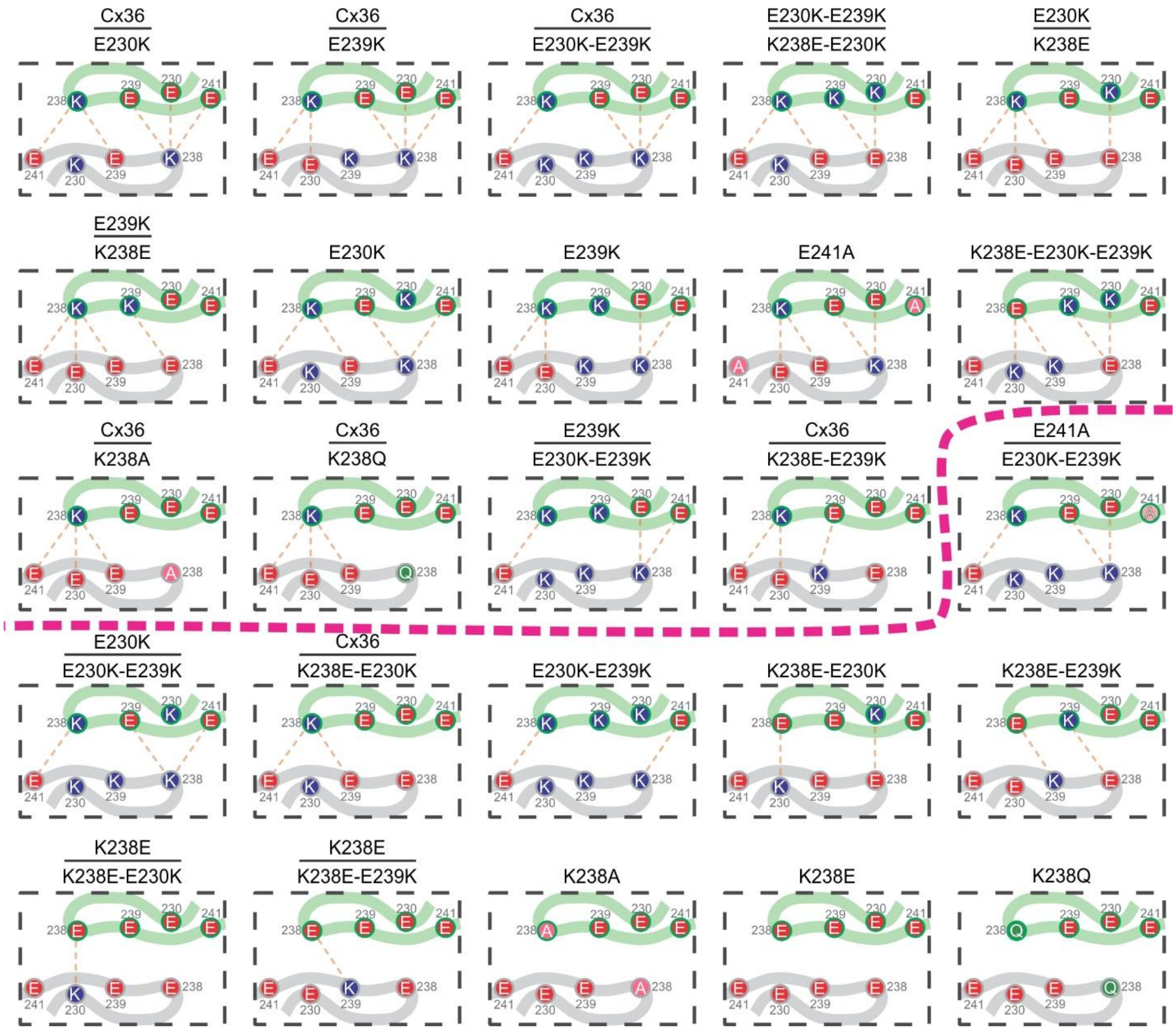


**Figure S2. Schematic of Cx36 E2-E2 docking interface and predicted ESIs for each variant combination.** A portion of docked E2-E2 domains from apposing connexins is colored in green and grey. Orange dashed lines indicate the ESIs predicted to form at the E2-E2 docking interface between the indicated residues. The thick dashed pink line separates the functional variant GJ combinations (above) from the non-functional variant combinations (below) according to patch clamp data shown in Fig. 4.

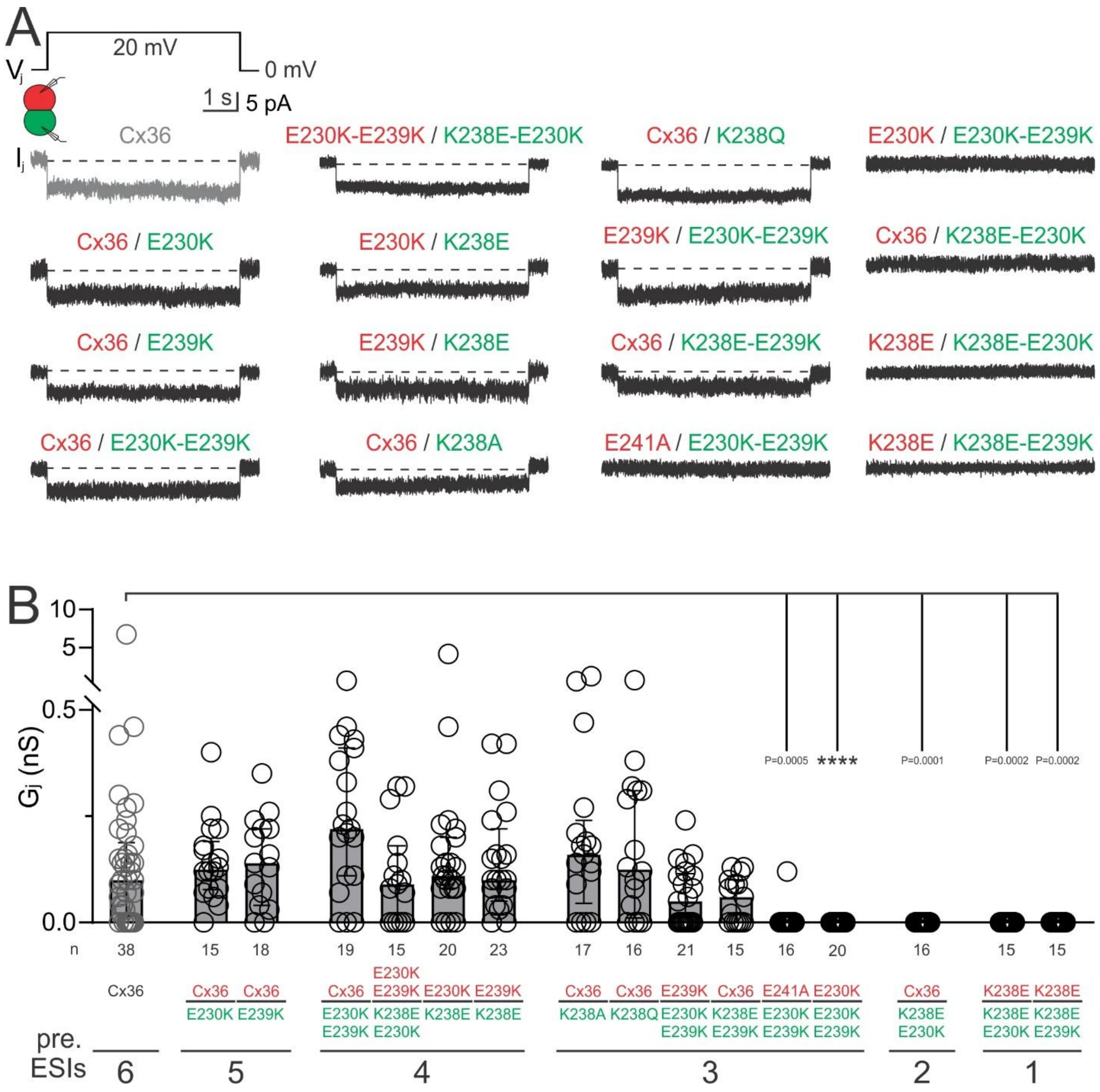


**Figure S3. Functional status of heterotypic Cx36 variant GJ combinations with various predicted numbers of ESIs at the E2-E2 docking interface.** (A) Representative $I_j$s were recorded in response to a $V_j$ pulse applied to DKO HEK293 cell pairs expressing a Cx36 variant-IRES-DsRed in one cell, and a different variant-IRES-EGFP in the other cell, as indicated by the colour of the text. (B) Bar graph summarizing the median $G_j$ (error bars represent IQR) for all cell pairs recorded. The number of predicted ESIs per E2-E2 is indicated for each group below the X-axis (pre. ESIs). A Kruskal-Wallis test followed by Dunn's multiple comparisons tests were performed to compare the $G_j$ of each Cx36 variant to that of Cx36 wildtype. Statistical significance is indicated on the graph (****$P < 0.0001$). "n" represents the number of biologically independent cell pairs measured for each group as indicated. $I_j$ and $G_j$s for Cx36 are shown here (same as Fig. 1) in grey for comparison purposes.

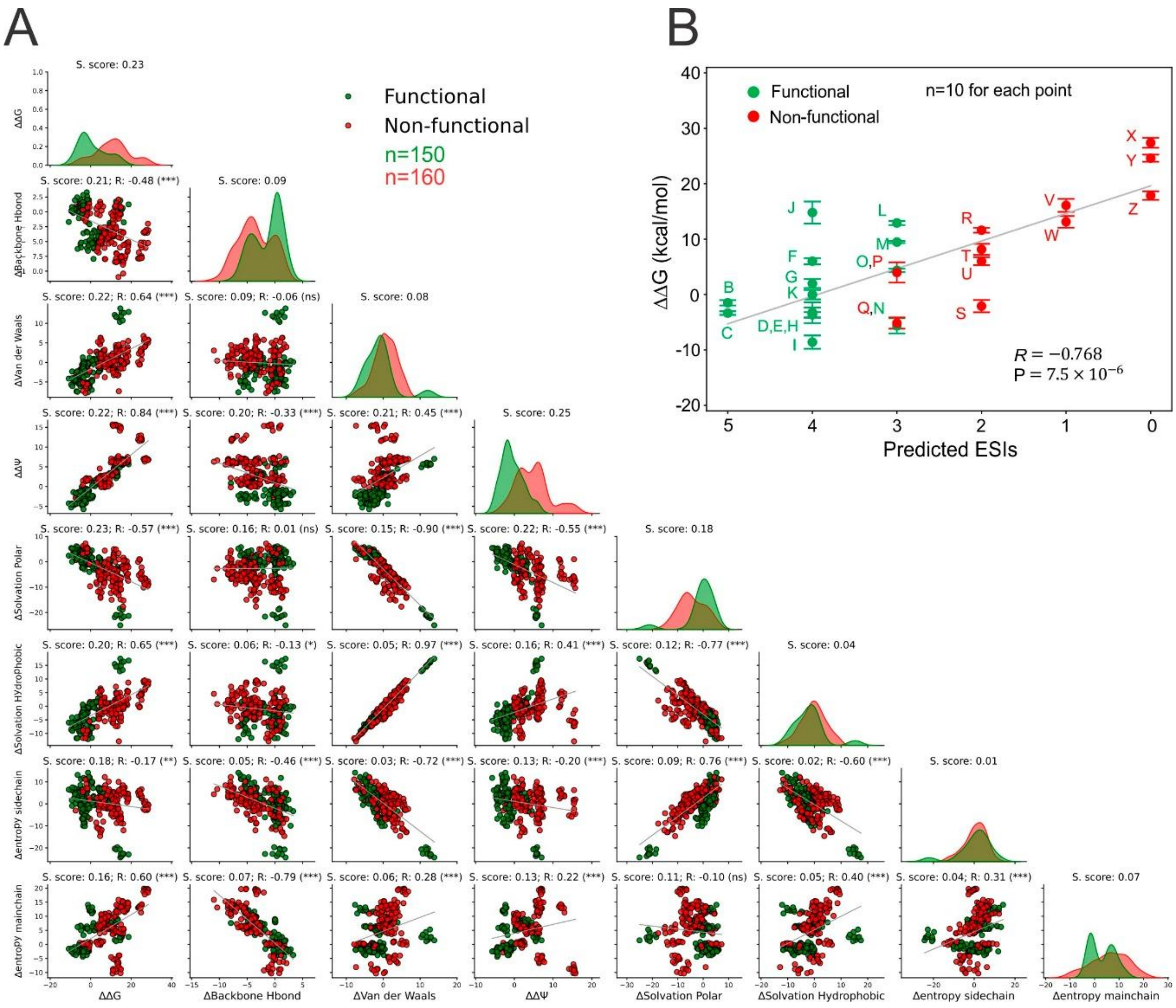


**Figure S4. Paired analysis of ΔΔG energy contributors assessed by Silhouette scores (S.score) and linear correlation between the predicted ESIs and ΔΔG for Cx36 GJ structure 8XGD.** (A) The diagonal panels display one-dimensional distribution plots for each individual energy term, where the Y-axis represents sample frequency. Off-diagonal panels show two-dimensional scatter plots for pairs of energy terms, illustrating their combined ability to separate functional (green dots) and non-functional (red dots) variant GJs. n denotes the sample size of the functional and non-functional distributions, obtained from 10 independent runs of calculation across 31 homo- and heterotypic variants. The separation quality is quantified by the Silhouette score (S. score on top of each plot). Pearson correlation coefficients (R) and statistical significance (ns, not significant; *P < 0.05; **P < 0.01; ***P < 0.001) are also indicated between each pair of energy terms. The unit for energy terms is kcal/mol. (B) ΔΔG represents the change in binding free energy of each variant GJ combination relative to the wild-type Cx36 GJ (i.e., mutant minus wild-type). Each variant GJ is labelled according to the corresponding letter as shown in Fig. 4A. n represents the independent runs of energy calculation for each variant. Error bars represent the computational standard deviation among 10 different runs. The Pearson correlation coefficient (R) and P-value indicate a statistically significant linear relationship between ESIs and ΔΔG.

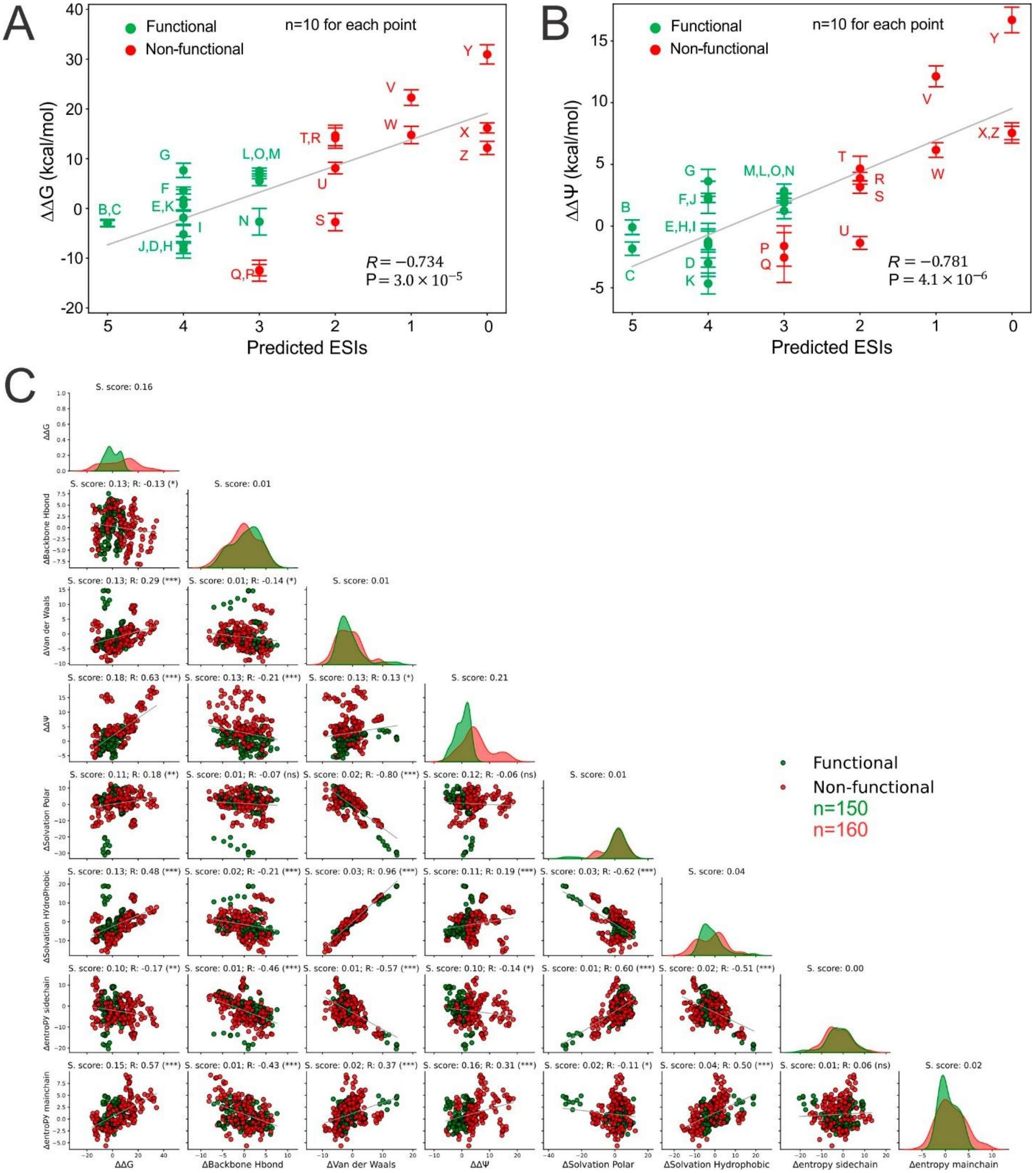


**Figure S5. Correlations of the predicted ESIs and ΔΔG/ΔΔΨ and paired analysis of ΔΔG energy contributors assessed by Silhouette scores (S.score) for Cx36 GJ structure 8IYG.** (A/B) ΔΔG/ΔΔΨ represents the change of binding free energy/electrostatics of the variant combination GJs relative to the wild-type Cx36 GJ (i.e., mutant minus wild-type). Each variant GJ is labelled according to Fig. 4A for successful (green) or unsuccessful (red) formation of functional GJs. n represents the independent runs of energy calculation for each variant. Error bars represent the standard deviation among 10 different runs. The Pearson correlation coefficient (R) and P-value indicate a statistically significant linear relationship between ESIs and ΔΔG/ΔΔΨ. (C) The diagonal panels display one-dimensional distribution plots for each individual energy term, where the Y-axis represents sample frequency. Off-diagonal panels show two-dimensional scatter plots for pairs of energy terms, illustrating their combined ability to separate functional (cyan dots) and non-functional (red dots) variant GJs. n denotes the sample size of the functional and non-functional distributions, obtained from 10 independent runs of calculation across 31 homo- and heterotypic variants. The separation quality is quantified by the Silhouette score (S. score on top of each plot). Pearson correlation coefficients (R) and statistical significance are also indicated between each pair of energy terms. The unit for energy terms is kcal/mol.

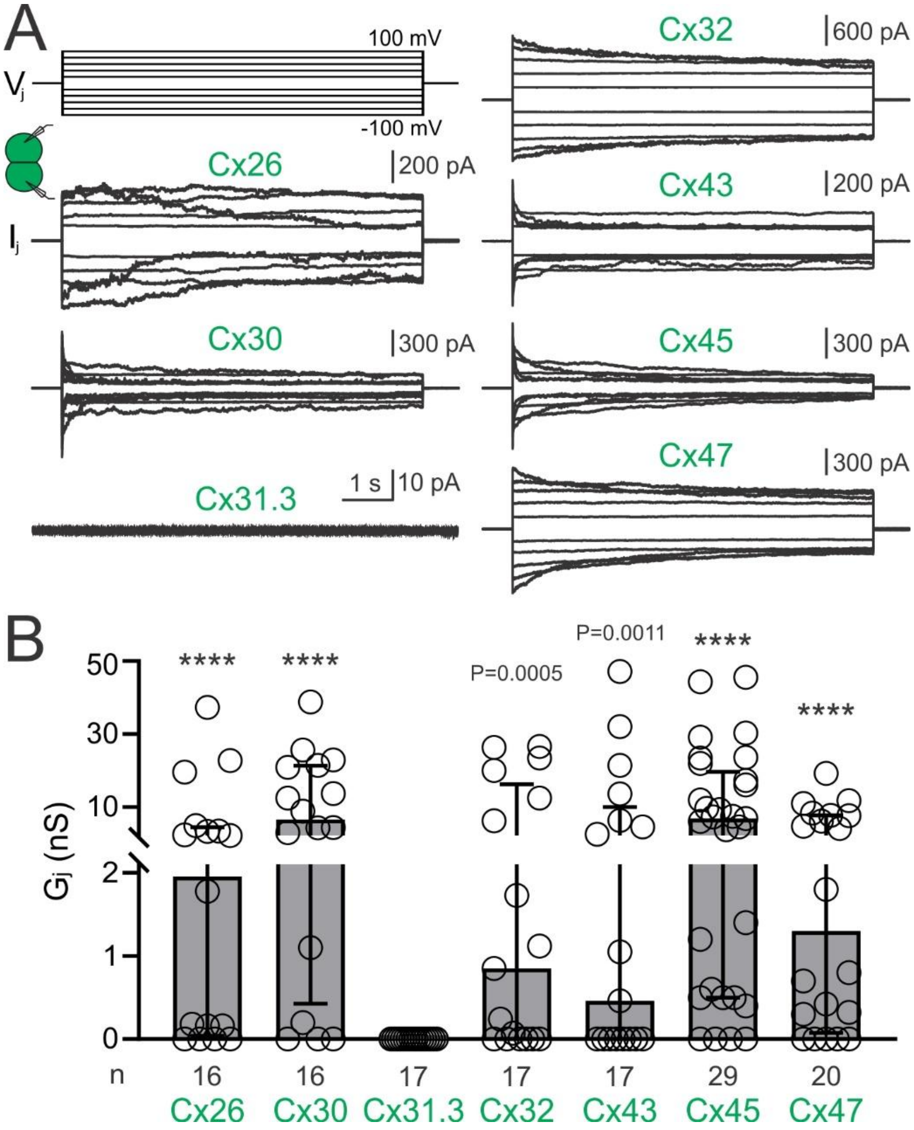


**Figure S6. Functional status of homotypic Cx26, Cx30, Cx31.3, Cx32, Cx43, Cx45, and Cx47 GJ channels.** (A) Superimposed $I_j$s were recorded in response to a series of $V_j$ pulses ranging from ±20 mV to ±100 mV applied to DKO HEK293 cell pairs expressing the indicated connexin. Expressing each of these connexins successfully formed functional GJs except Cx31.3. Signature $V_j$-gating properties were observed as expected for each of the connexins forming functional GJs. (B) Bar graph summarizing the median $G_j$ (error bars represent IQR) for all cell pairs recorded. Mann-Whitney tests were performed to compare the $G_j$ of each homotypic connexin GJ to their respective heterotypic GJ formed with Cx36 (shown in Fig. 5), where statistical significance is indicated above the respective homotypic GJ as indicated. Cx31.3 failed to form any functional GJs and showed no statistical difference from the $G_j$ of Cx31.3 / Cx36 GJs. "n" represents the number of biologically independent cell pairs measured for each group as indicated.The replating time prior to patch clamp for Cx32 and Cx43 homotypic GJs was 30 minutes, and for other connexins was 2 hours. Shorter replating time for Cx32 and Cx43 was due to the fact that these connexins tend to form too high GJ coupling with $G_j$ in a range indistinguishable from apparent cell pairs not yet finished cell division (also known as mitotic bridge). The caveats of shorter replating time were an underestimation of $G_j$ and an increased number of false negatives in these cell pairs.

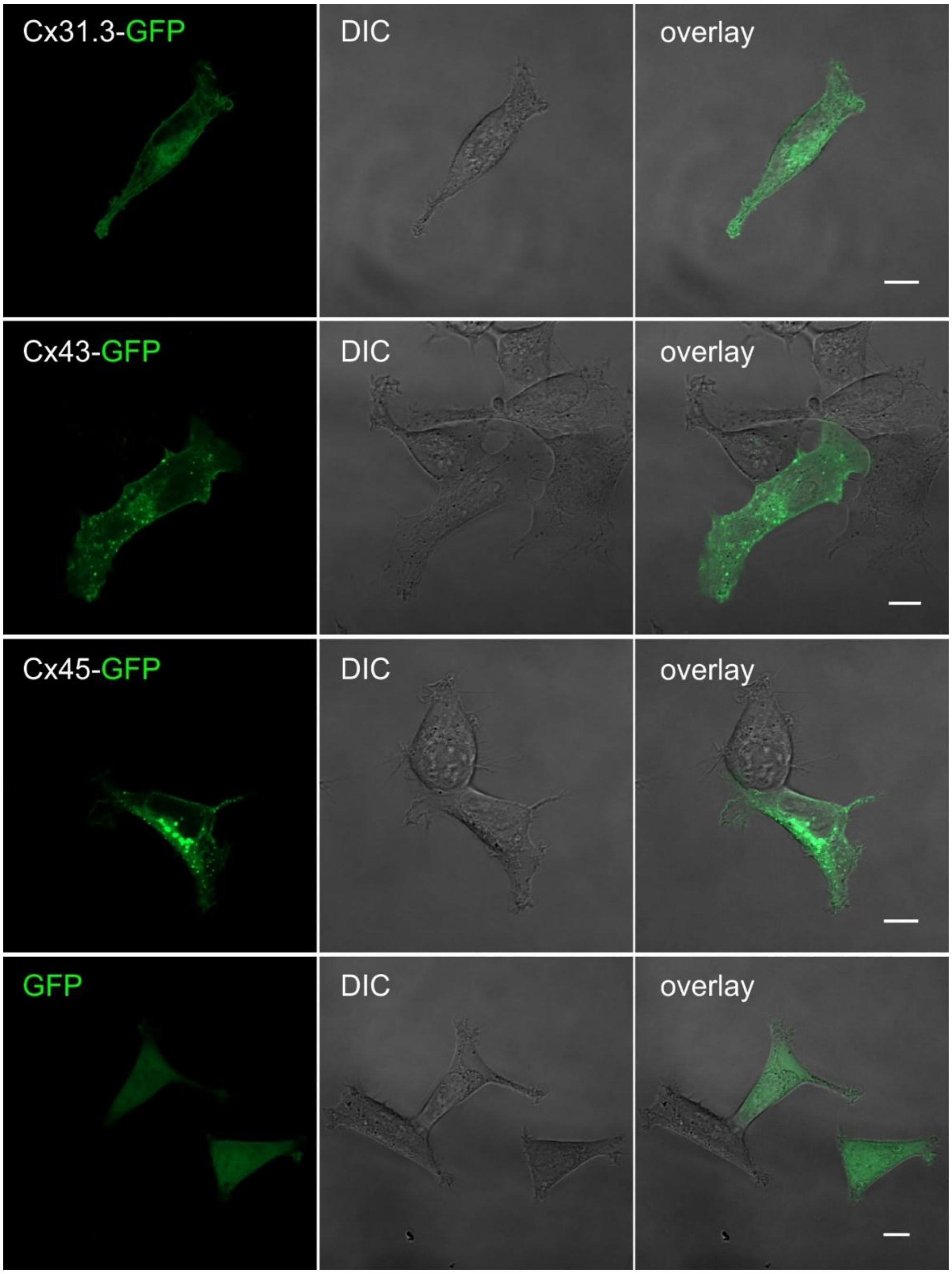


**Figure S7. Localization of GFP-tagged Cx31.3, Cx43, Cx45 in DKO HEK293 cells.** Fluorescent (left panels), differential interference contrast (DIC, middle panels) and superimposed (overlay, right panels) images are shown for DKO HEK293 cells expressing carboxyl-terminal tagged connexins as indicated, Cx31.3-GFP, Cx43-GFP, and Cx45-GFP. Cells expressing GFP alone were used as a control. Cx31.3-GFP typically showed a localization around the nuclei and apparently on the plasma membrane, similar to those observed for Cx43-GFP and Cx45-GFP. However, GFP is evenly distributed throughout the nuclei and cytosol without any preferential localization. Scale bars = 10 μm.

**Functional**

| Variant GJs | ESIs | Label |
|---|---|---|
| Cx36 | 6 | A |
| Cx36 / E230K | 5 | B |
| Cx36 / E239K | | C |
| Cx36 / E230K-E239K | 4 | D |
| E230K-E239K / K238E-E230K | | E |
| E230K / K238E | | F |
| E239K / K238E | | G |
| E230K | | H |
| E239K | | I |
| E241A | | J |
| K238E-E230K-E239K | | K |
| Cx36 / K238A | 3 | L |
| Cx36 / K238Q | | M |
| E239K / E230K-E239K | | N |
| Cx36 / K238E-E239K | | O |

**Non-Functional**

| Variant GJs | ESIs | Label |
|---|---|---|
| E241A / E230K-E239K | 3 | P |
| E230K / E230K-E239K | | Q |
| Cx36 / K238E-E230K | 2 | R |
| E230K-E239K | | S |
| K238E-E230K | | T |
| K238E-E239K | | U |
| K238E / K238E-E230K | 1 | V |
| K238E / K238E-E239K | | W |
| K238A | 0 | X |
| K238E | | Y |
| K238Q | | Z |

**Figure S8. Non-functional variants were able to form functional GJs in different combinations.** Table of functional and non-functional Cx36 variants tested. Boxes and arrows of the same colour highlight the same variant tested in different non-functional and functional combinations. Each non-functional variant was shown in at least one other combination to have the ability to form functional GJs, demonstrating their ability to localize to the cell surface.
